# Functional network collapse in neurodegenerative disease

**DOI:** 10.1101/2023.12.01.569654

**Authors:** Jesse A. Brown, Alex J. Lee, Kristen Fernhoff, Taylor Pistone, Lorenzo Pasquini, Amy B. Wise, Adam M. Staffaroni, Maria Luisa Mandelli, Suzee E. Lee, Adam L. Boxer, Katherine P. Rankin, Gil D. Rabinovici, Maria Luisa Gorno Tempini, Howard J. Rosen, Joel H. Kramer, Bruce L. Miller, William W. Seeley, Alzheimer’s Disease Neuroimaging Initiative (ADNI)

## Abstract

Cognitive and behavioral deficits in Alzheimer’s disease (AD) and frontotemporal dementia (FTD) result from brain atrophy and altered functional connectivity. However, it is unclear how atrophy relates to functional connectivity disruptions across dementia subtypes and stages. We addressed this question using structural and functional MRI from 221 patients with AD (n=82), behavioral variant FTD (n=41), corticobasal syndrome (n=27), nonfluent (n=34) and semantic (n=37) variant primary progressive aphasia, and 100 cognitively normal individuals. Using partial least squares regression, we identified three principal structure-function components. The first component showed overall atrophy correlating with primary cortical hypo-connectivity and subcortical/association cortical hyper-connectivity. Components two and three linked focal syndrome-specific atrophy to peri-lesional hypo-connectivity and distal hyper-connectivity. Structural and functional component scores predicted global and domain-specific cognitive deficits. Anatomically, functional connectivity changes reflected alterations in specific brain activity gradients. Eigenmode analysis identified temporal phase and amplitude collapse as an explanation for atrophy-driven functional connectivity changes.

## Introduction

Cognitive deficits in Alzheimer-type dementia (AD) and frontotemporal dementia (FTD) result from tissue degeneration in specific brain regions ^1–11^. Brain functional connectivity alterations are also common, involving both decreases and increases in connectivity when compared to cognitively unimpaired individuals ^12–18^. A major challenge in clinical neuroscience is to understand the relationship between structural and functional alterations and what overlapping or unique contributions they make to cognitive impairment ^19–23^. Progress on this question requires datasets and methods that can unravel the subtypes and stages of atrophy ^24^ and map these to distinct patterns of functional hypo and hyper-connectivity ^25^.

Recent advances in functional brain activity modeling may help explain structure-function relationships in neurodegenerative disease. Canonical functional networks are embedded in a small number of spatial gradients that can be identified with dimensionality reduction techniques ^26,27^. These gradients appear to represent intrinsic systems that comprise a low-dimensional basis for different activity and connectivity states ^28–30^. In the current study, we hypothesized that atrophy-associated functional connectivity alterations could be parsimoniously explained by disruptions of specific spatial gradients. This endeavor required reconciling two key sets of findings. The first set of findings is that patients with AD and behavioral variant FTD (bvFTD) exhibit opposing spatial patterns of hypo and hyper-connectivity ^14,31^. AD involves posterior-predominant atrophy and functional connectivity reductions in the default mode network. By contrast, bvFTD involves fronto-insular atrophy and connectivity reductions in the salience network ^12,32^. The second key set of findings is that patients with Parkinson’s disease or AD have weaker functional connectivity in primary sensory networks – regions that are remote from the primary sites of pathology and neurodegeneration – and stronger connectivity in subcortical and/or association networks ^33,34^. Thus, at least two different types of atrophy-associated connectivity alteration are apparent: 1) hypo-connectivity near the lesion and hyper-connectivity remote from it; 2) a convergent pattern of sensory network hypo-connectivity and association network hyper-connectivity across syndromes with different atrophy patterns. Intriguingly, anti-correlated networks are unified as opposing poles of individual gradients ^30,35^, raising the possibility that different atrophy patterns may disrupt distinct gradients and cause hypo and hyper-connectivity as two sides of the same coin.

Here we studied structure-function relationships using a rich dataset of structural and functional MRI scans from 221 patients with Alzheimer’s-type dementia, behavioral variant FTD, corticobasal syndrome (CBS), nonfluent and semantic variants of primary progressive aphasia (nfvPPA/svPPA), and 100 age-matched cognitively normal (CN) controls subjects. We identified three principal structure-function components linking different atrophy patterns to specific brain-wide functional connectivity alterations.

These structural and functional components made independent contributions to cognitive deficits. Our analysis revealed that functional connectivity decreases and increases were linked to alterations in a small set of intrinsic functional gradients. Specifically, we found evidence that atrophy associates with reductions in gradient amplitude and changes in between-gradient phase coupling. These two processes reflect both the common and distinct patterns of connectivity alteration across patients with different atrophy subtypes and stages.

## Results

### Functional connectivity decreases and increases across the atrophy spectrum

We assessed structure-function relationships in 221 patients across the FTD-AD spectrum and 100 age, sex, scanner, and fMRI head motion-matched cognitively normal controls, comprising the study’s primary cohort (**Table 1**). We measured gray matter atrophy in 246 cortical and subcortical regions using regional W-scores (**Methods**). Task-free functional connectivity (FC) was measured between region pairs. 97% of regions had significant gray matter atrophy (W > 1.5) in at least five subjects, showing that these clinical syndromes collectively involve the entire brain. We used partial least squares regression (PLSR) to identify the primary structure-function components. The first three structure components had high reproducibility (**Methods** and **Supplementary Figure 1**) and were made the focus of this study.

**Table 1.**
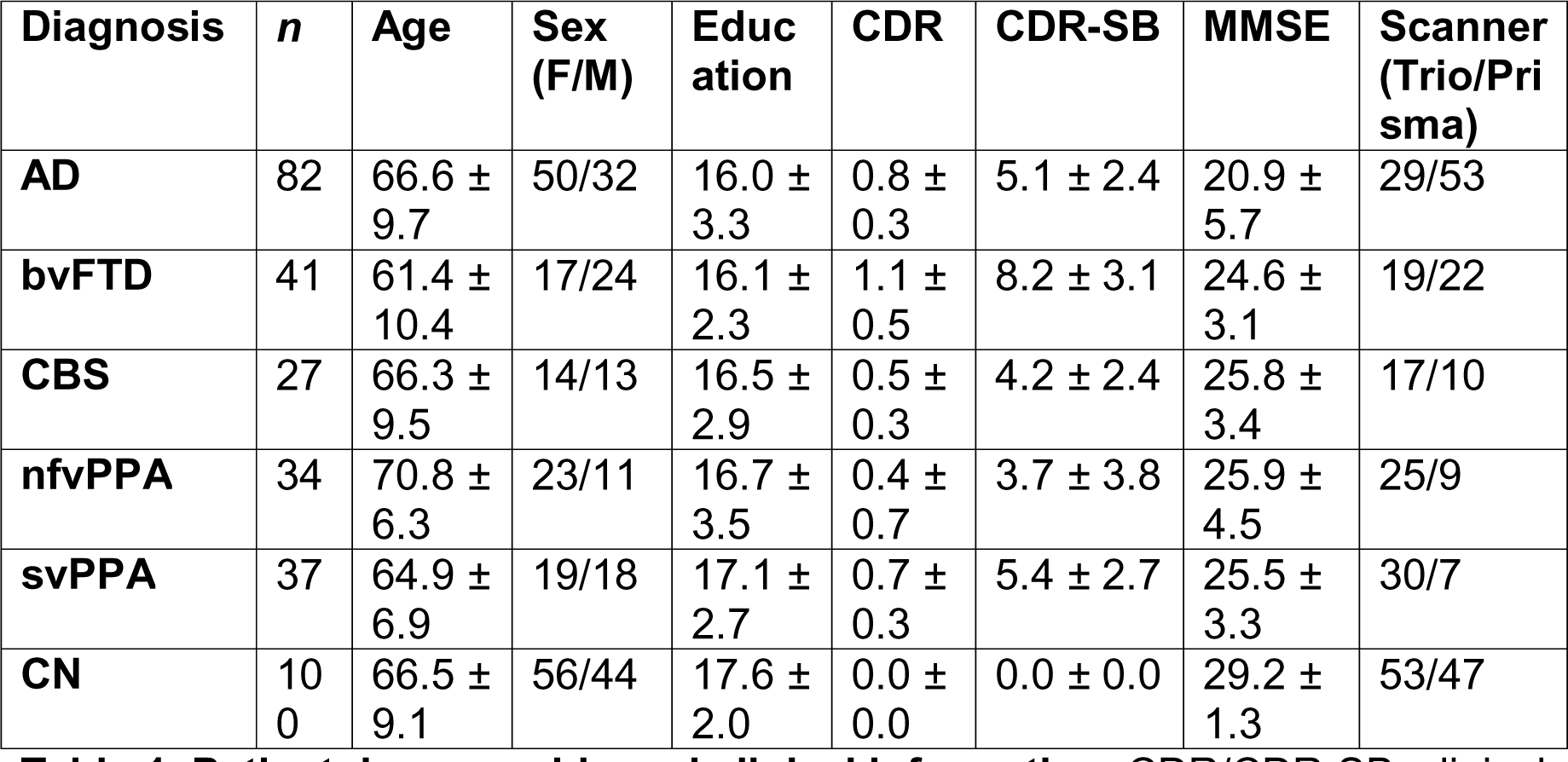
Patient demographic and clinical information. CDR/CDR-SB: clinical dementia rating/sum of boxes; MMSE: mini-mental status examination.

| Diagnosis | *n* | Age | Sex (F/M) | Education | CDR | CDR-SB | MMSE | Scanner (Trio/Prisma) |
| --- | --- | --- | --- | --- | --- | --- | --- | --- |
| AD | 82 | 66.6 ± 9.7 | 50/32 | 16.0 ± 3.3 | 0.8 ± 0.3 | 5.1 ± 2.4 | 20.9 ± 5.7 | 29/53 |
| bvFTD | 41 | 61.4 ± 10.4 | 17/24 | 16.1 ± 2.3 | 1.1 ± 0.5 | 8.2 ± 3.1 | 24.6 ± 3.1 | 19/22 |
| CBS | 27 | 66.3 ± 9.5 | 14/13 | 16.5 ± 2.9 | 0.5 ± 0.3 | 4.2 ± 2.4 | 25.8 ± 3.4 | 17/10 |
| nfvPPA | 34 | 70.8 ± 6.3 | 23/11 | 16.7 ± 3.5 | 0.4 ± 0.7 | 3.7 ± 3.8 | 25.9 ± 4.5 | 25/9 |
| svPPA | 37 | 64.9 ± 6.9 | 19/18 | 17.1 ± 2.7 | 0.7 ± 0.3 | 5.4 ± 2.7 | 25.5 ± 3.3 | 30/7 |
| CN | 100 | 66.5 ± 9.1 | 56/44 | 17.6 ± 2.0 | 0.0 ± 0.0 | 0.0 ± 0.0 | 29.2 ± 1.3 | 53/47 |

The first structure-function component (SF1) accounted for 51.2% of the brain atrophy variance and captured the relationship between overall mean atrophy and a distributed pattern of FC decreases and increases. For each component, each subject received structure and function scores. This component’s structure scores correlated perfectly with subject overall mean atrophy (r=0.994). The SF1 structure-function score correlation was r=0.56, p < 0.001 (**Figure 1A**, left). The FC pattern associated included more negative FC between primary visual, somatomotor, and auditory regions (**Figure 1B**, left). There was more positive subcortical-cortical FC, most strongly between the striatum, thalamus, and motor regions, along with more subtly increased fronto-parietal association cortex connectivity to widespread cortical and subcortical regions. This pattern captured statistically significant differences in FC edge strength between subjects with low/medium/high scores on structure component 1 (**Supplementary Figure 2**).

**Figure 1.**
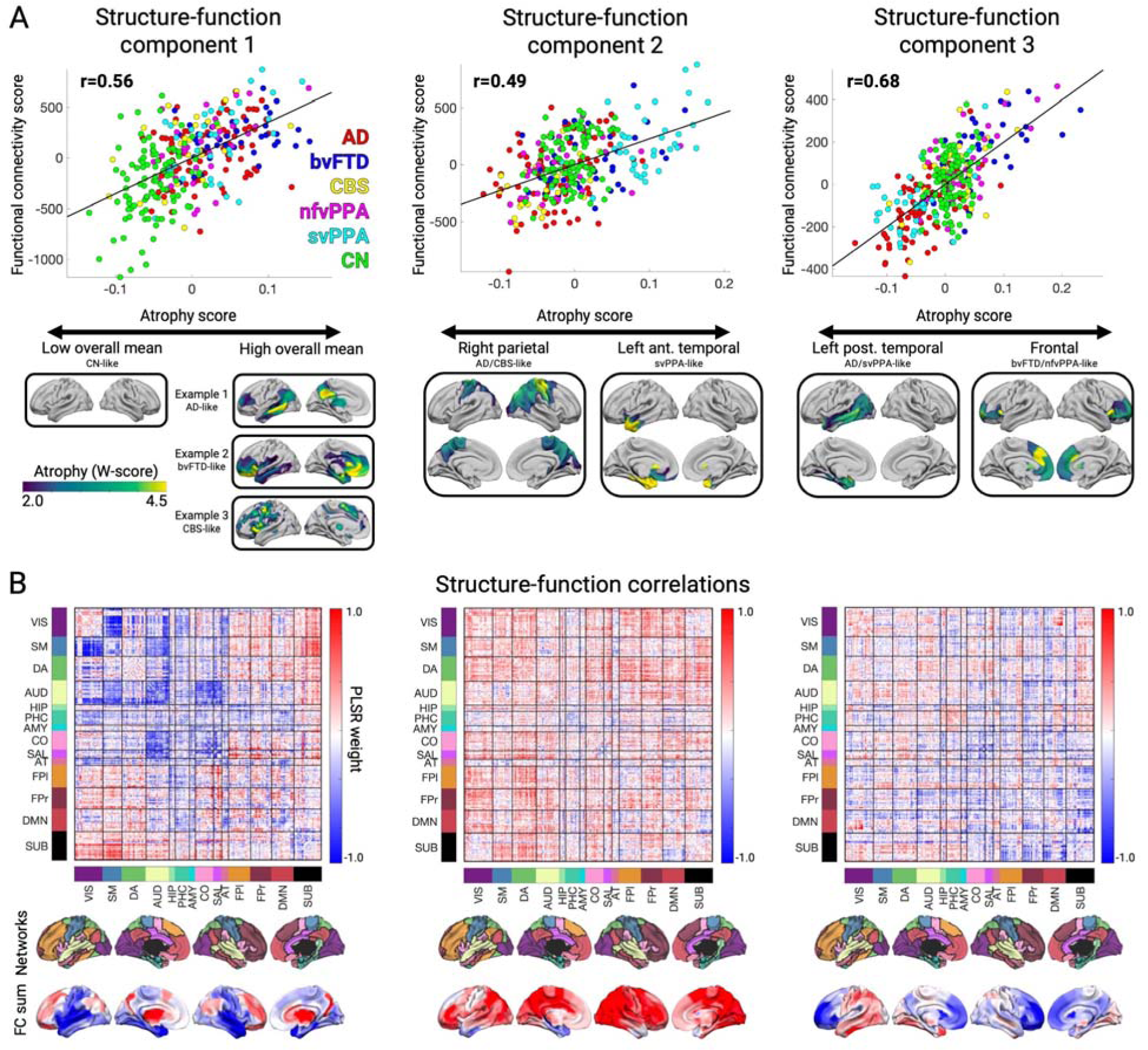
The first three structure-function components across the AD-FTD spectrum. **A.** The correlation between atrophy and functional connectivity (FC) scores for partial least squares regression components 1-3. Beneath each correlation plot are the associated atrophy patterns associated with a negative or positive atrophy component score. For component 1, three example patients are shown with different atrophy patterns but equivalent overall mean atrophy. **B.** Matrices showing the partial least squares FC weights for each component, along with the network membership for each brain region. Negative and positive weights indicate decrease or increase in FC with an increase in the atrophy component score (and vice versa). Matrices show 14 brain FC networks for reference. The top row of brain surfaces shows the 14 networks. The bottom row of brain surfaces shows the FC PLSR region sums. VIS: visual; SM: sensory-motor; DA: dorsal attention network; AUD: auditory; HIP: hippocampal; PHC: parahippocampal; AMY: amygdala; CO: cingulo-opercular; SAL: salience; AT: anterior temporal; FPl: left fronto-parietal; DMN: default mode network; FPr: right fronto-parietal; SUB: subcortical.

The second and third components captured syndrome-specific atrophy patterns that explained 9.1% and 6.5% of the atrophy variance. The structure-function correlation on Component 2 was r=0.49, p < 0.001; (Figure 1A**, middle**). Patients on the positive end of the Component 2 spectrum had svPPA-like atrophy in the left anterior temporal lobe. This accompanied weakened functional connectivity (lower than controls) from anterior and medial temporal regions, both locally and globally (Figure 1B**, middle**).

These patients showed heightened functional connectivity involving the dorsal attention, visual, and fronto-parietal networks. In contrast, patients at the negative end of the Component 2 spectrum had AD or CBS diagnoses and had the opposite pattern of atrophy and functional connectivity: atrophy in the right dorsal parietal cortex, sensory-motor cortex, and thalamic areas; peri-atrophy connectivity deficits; and elevated FC in the left anterior temporal lobe. Cognitively normal control subjects appeared in the middle of the spectrum, with minimal brain atrophy and balanced FC in anterior temporal and dorsal parietal-anchored networks. The Component 3 structure-function correlation was r=0.68, p < 0.001 (Figure 1A**, right**). Subjects with low/medium/high scores on atrophy components 2 and 3 also had progressive differences in overall FC edge strength (**Supplementary Figure 2**).

We used the three structural components to assess syndrome differences in atrophy and functional connectivity. First, we reduced the three atrophy components to two dimensions using multidimensional scaling to visualize the AD-FTD atrophy spectrum (Figure 2). We then identified the subset of patients for each syndrome expressing the “typical” atrophy pattern and the corresponding functional connectivity alterations. AD patients showed atrophy in the posterior temporal and parietal lobe; bvFTD in the insula, rostral/orbital frontal lobe, and anterior cingulate; CBS in the primary sensory/motor cortex and superior frontal lobe; nfvPPA in the inferior frontal lobe, insula, and premotor cortex; and svPPA in the anterior temporal lobe. The associated functional connectivity alterations are shown in Figure 2. We then tested how well those FC patterns could be explained in terms of the three functional components. For each syndrome, the best match to the true FC difference matrix (Figure 2, upper triangles) was the corresponding reconstructed FC matrix (Figure 2, lower triangles) (AD, actual versus reconstructed r=0.81; bvFTD r=0.82; CBS r=0.41; nfvPPA r=0.50; svPPA r=0.82). This confirmed that three structure-function components captured the predominant syndrome-associated patterns.

**Figure 2.**
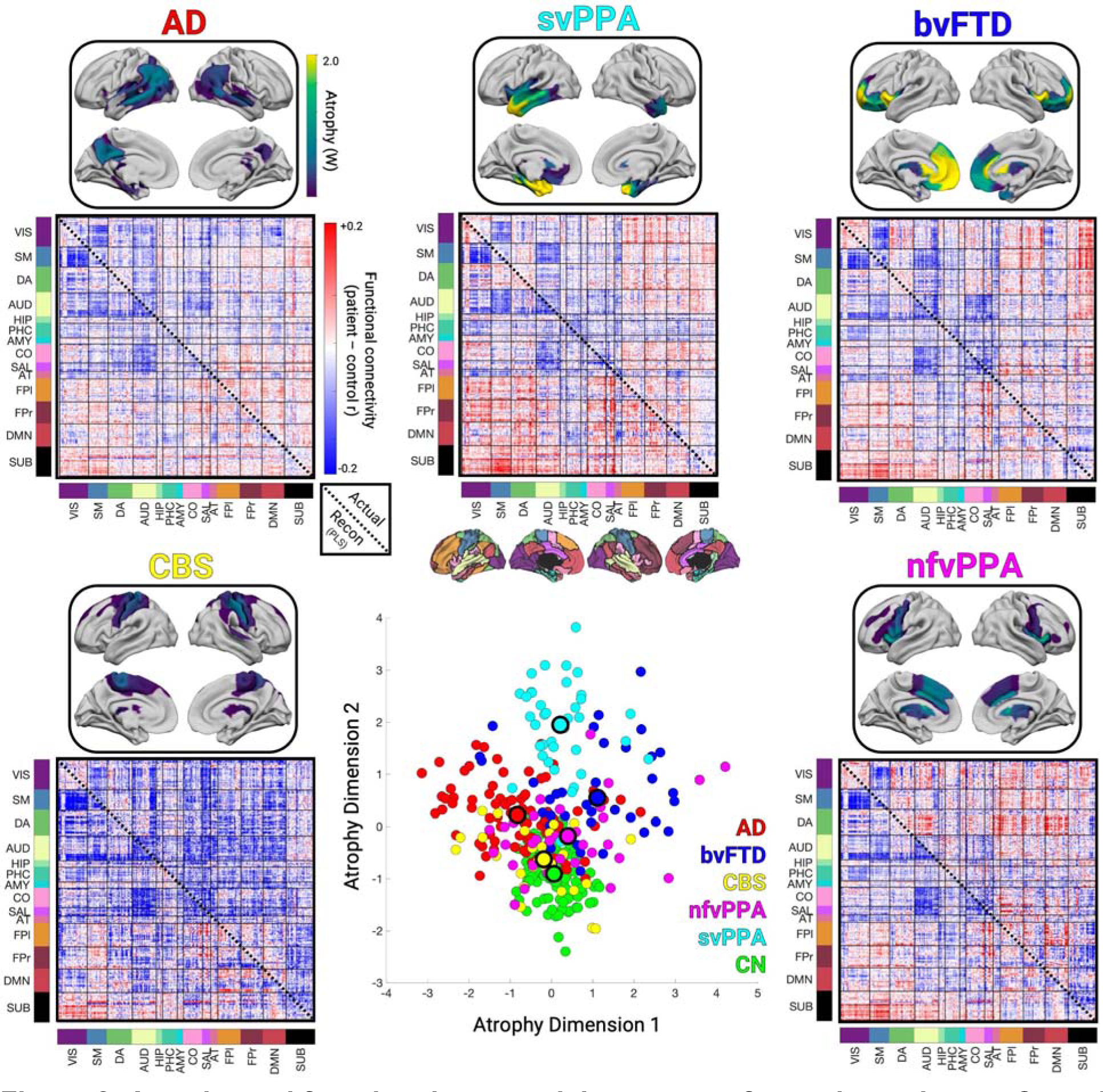
Atrophy and functional connectivity patterns for each syndrome. Central scatter plot depicts individual subjects’ atrophy similarity in two dimensions based on multidimensional scaling of structure components 1-3. Large dots show the mean position for each clinical syndrome. For each syndrome, the mean atrophy pattern and functional connectivity matrix are shown for the set of “typical” patients that were accurately classified as having that syndrome based on their individual atrophy pattern. The mean functional connectivity difference matrix is shown for the typical patients for each syndrome versus cognitively normal subjects (upper triangles), along with the reconstructed matrix based on function components 1-3 (lower triangles).

We confirmed the structure-function relationship reliability in two ways. First, we used ridge regression with four-fold cross-validation (**Methods**). We found that the three primary components each had significant structure-function correlation in the left-out fold (SF1: median r=0.49, median p < 0.001; SF2: r=0.32, p=0.003; SF3: r=0.39, p < 0.001; **Supplementary Figure 3A**). Importantly, the FC edge weights associated with each atrophy component in cross-validated subsamples strongly matched the FC edge weights for the corresponding component from the full sample (**Supplementary Figure 3B**). Second, we assessed these structure-function components in a separate replication dataset from the Alzheimer’s Disease Neuroimaging Initiative (ADNI3) sample, comprised of 421 cognitively normal subjects and 56 subjects with Alzheimer’s disease from 35 scanning sites. Structure and function scores for each component were significantly correlated (SF1: partial r=0.25, t=4.94, p < 0.001; SF2: partial r=0.15, t=3.81, p < 0.001; SF3: partial r=0.08, t=2.42, p=0.015; **Supplementary Figure 4**). Collectively, the three main structure-function components in AD and FTD captured specific atrophy patterns and associated hypo/hyper-FC profiles that replicated in different individuals, syndromes, and MRI scanners.

### Neuropsychological correlations with structural and functional components

We next examined the relationship between the brain structure-function components and cognitive performance. We first focused on two tests of global functioning, the CDR®+NACC-FTLD sum of boxes (Miyagawa et al., 2020; henceforth referred to as CDR-SB) and MMSE. A generalized additive model was used to estimate cognitive scores based on the first three structure and function scores, either as linear or non-linear terms, and covariates (**Methods**). The model for CDR-SB explained 50% of the variance. The strongest predictors were S1 (F=22.88, p < 0.001; Figure 3A**/B**), F1 (F=17.16, p < 0.001), and S3 (F=10.74, p < 0.001). This indicated that patients with the most severe clinical impairment had high overall mean atrophy (S1), most pronounced in the frontal lobe (S3), along with subcortical hyperconnectivity and primary sensory cortical hypoconnectivity (F1). For MMSE, the model explained 40% of the variance with the strongest predictions from S1 (F=22.31, p < 0.001), F3 (F=13.94, p < 0.001), and S3 (F=7.22, p=0.001). S3 had significant nonlinearity (Figure 3B), such that subjects with either frontal or temporal atrophy had equivalently poor MMSE scores. Subjects with worse MMSE scores also had anterior connectivity deficits and posterior connectivity enhancements. Thus, CDR-SB and MMSE scores significantly correlated with the overall amount of atrophy, specific atrophy patterns, and distinct functional connectivity alterations.

**Figure 3.**
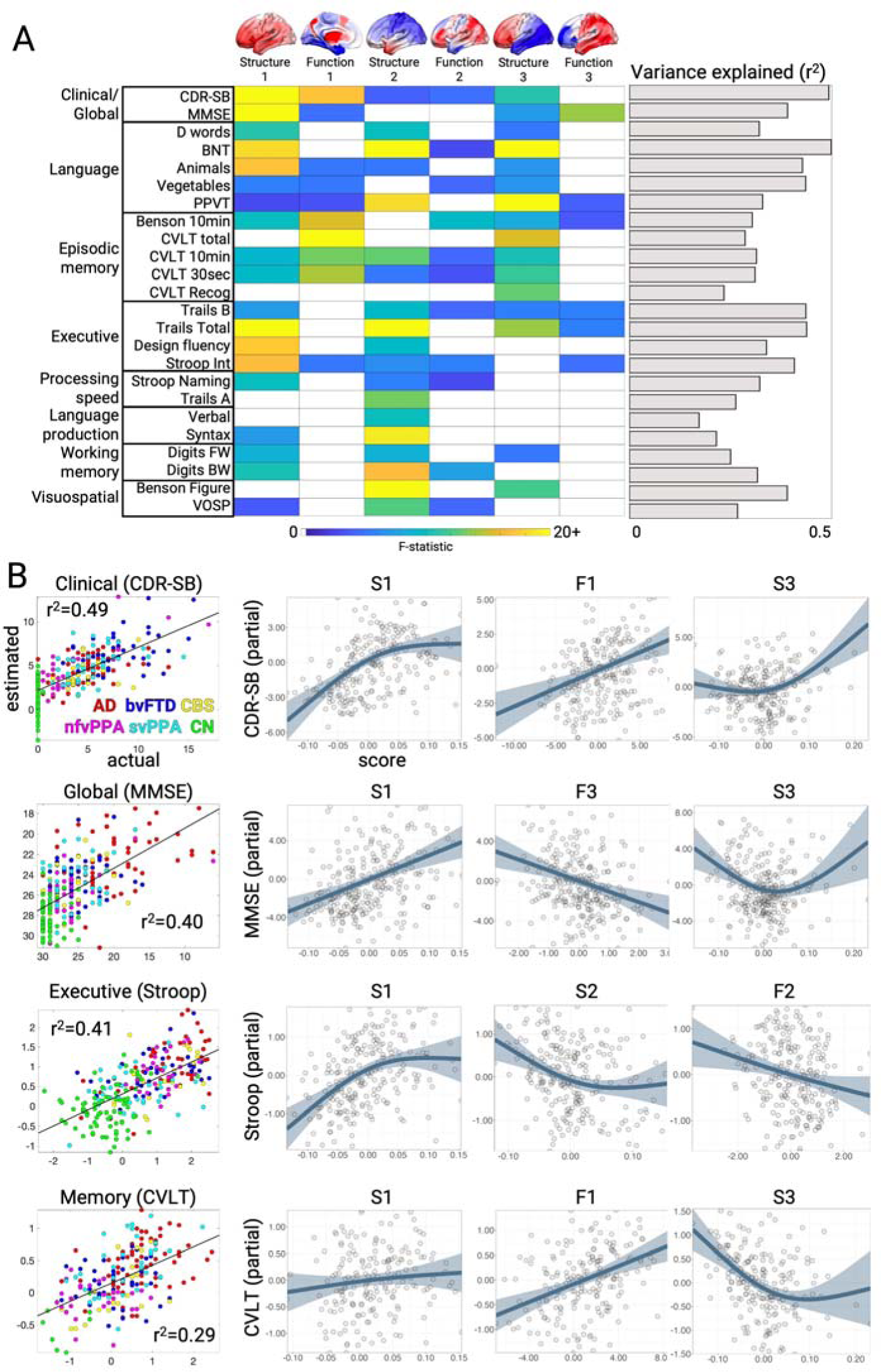
Neuropsychological correlates of brain structure-function components. **A.** Cognitive test score estimates based on a generalized additive model with brain structure-function component scores and covariates. The F-statistic for the structure and function terms in each model are shown when significant (p < 0.05, FDR-corrected) or trending (p < 0.05, uncorrected). The overall variance explained for each cognitive test is also shown. Brains show the weight of each region for structure components and the sum of region FC edge weights for function components. **B.** Correlation between actual and estimated test scores for clinical dementia severity (CDR-SB), global cognition (MMSE), executive functioning (Stroop Interference), and episodic memory (California Verbal Learning Test, CVLT total). Test scores are presented with higher values representing worse performance, and converted to Z-scores for Stroop and CVLT. Partial effect plots are shown for three predictors of interest for each test. Positive relationships indicate a correlation between the positive structure/function pattern and the neuropsychological score. Shaded bands show ± 2 x standard errors of the fit (95% confidence interval). BNT: Boston Naming Test; PPVT: Peabody Picture Vocabulary Test; VOSP: Visual Object and Space Perception.

We then assessed the relationship between brain structure-function component scores and neuropsychological test scores for episodic memory, working memory, processing speed, executive function, visuospatial processing, speech, and language. Across domains, the models explained an average of 34% of the variance (Figure 3A; mean r^2^=0.34±0.09; range=0.17-0.50). The brain-behavior relationships clustered into two groups. The first group represented dysfunctions in clinical/global functioning, language, and episodic memory, most strongly influenced by S1, F1, and S3. This pattern was most strongly exhibited by patients with AD or svPPA, with high mean atrophy, most prominent in the temporal lobe, along with sensory hypo-connectivity and subcortical/association hyper-connectivity. The second group included dysfunctions in executive function, processing speed, language production, and visuospatial processing, driven by S1 and S2. The patients most strongly represented in this group had CBS, nfvPPA, or AD, also with high mean atrophy but focused in the parietal lobe. Overall, brain-behavior relationships were strongest for structural components, though key functional relationships predicted global cognition and memory performance.

We evaluated the reliability of the structure-function-cognition relationship in two ways. First, we estimated CDR-SB scores in the ADNI replication dataset using 774 scans with associated cognitive test scores. This model explained 33.2% of the CDR-SB variance with the strongest predictions from S1 (F=57.99, p < 0.001), F1 (F=12.13, p < 0.001), F3 (F=21.33, p < 0.001), and S2 (F=16.63, p < 0.001). This indicated that the same structure-function patterns were present in different patients with dementia and had a largely similar impact on cognitive impairment. Second, we evaluated the longitudinal relationship between S1, F1, and CDR-SB using baseline and follow-up data from a subset of 47 patients and 6 cognitively normal subjects from the main dataset (mean visit interval=1.1±0.5 years, range=0.4-2.6 years). We found that within-subject CDR-SB change significantly correlated with F1 change (t=2.05, p=0.04; **Supplementary Figure 5**) and S1 change (t=2.15, p=0.04). Between-subject CDR-SB mean was significantly related to S1 mean (F=6.06, p=0.02) and F1 mean (F=3.34, p=0.03) as expected from the cross-sectional model. A brain-only model showed that overall F1 correlated with both S1 mean (F=21.42, p < 0.001) and S1 change (F=6.21, p=0.02). Thus within-subject changes in structure and function component 1 were related, and both tracked a patient’s change in clinical dementia severity.

### Low-dimensional functional connectivity changes associate with different atrophy components

All three primary structure-function components involved hypo and hyper-FC. We hypothesized that specific alterations in low-dimensional brain activity underlie the hypo/hyper-FC patterns for each functional component. This was tested by performing PCA dimensionality reduction on the fMRI timeseries data to derive spatial components, henceforth referred to as gradients (Figure 4A), and their associated temporal fluctuations (**Methods**). Specifically, we derived the PCA space from an independent cohort of age-matched cognitively normal subjects (n=321) and projected all primary cohort patient and control fMRI data into this space. This approach assumed spatial gradient patterns are stable regardless of disease, and that atrophy perturbs gradient temporal dynamics. The spatial patterns captured known gradients including unipolar sensory-to-association (Gradient 1), sensory-to-cognitive (Gradient 2), visual-to-sensorimotor (Gradient 3), task-negative-to-task-positive (Gradient 4), and left-right asymmetric (Gradient 6) ^26,30^. We confirmed that the gradient spatial maps derived from the independent cohort were highly similar to those obtained from the primary cohort (**Supplementary Figure 6**).

**Figure 4.**
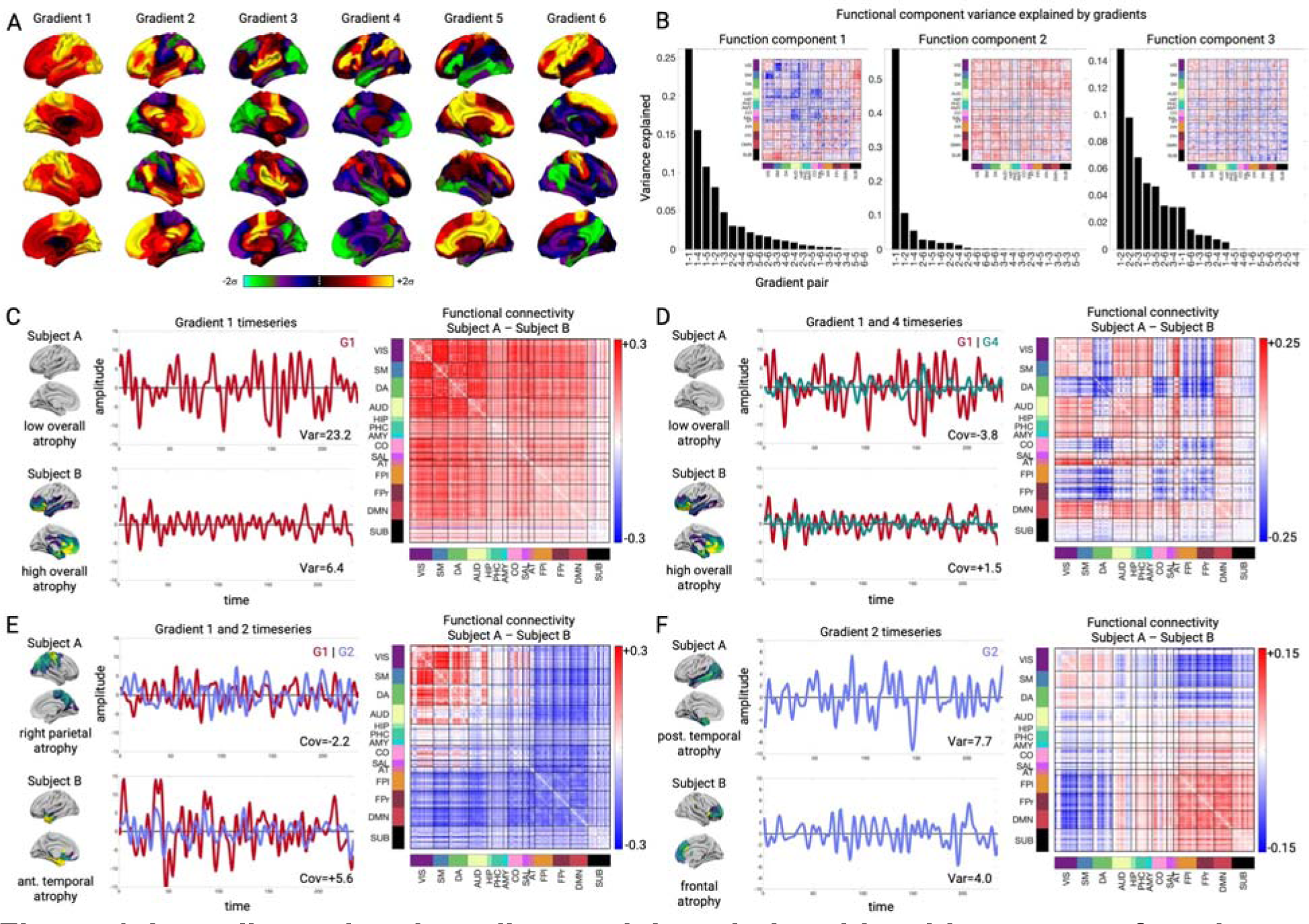
Low-dimensional gradient activity relationship with structure-function components. **A.** Gradient spatial weight maps based on PCA of fMRI timeseries data from the independent cognitively normal cohort (n=321). Weights represent PCA loadings. **B.** Variance explained in each PLSR functional component by individual gradient variances (e.g. 1-1, 2-2) and gradient pair covariances (e.g. 1-2, 2-3). Insets show PLSR component functional connectivity (FC) edge weight matrices. **C-F.** Select gradient timeseries and associated FC differences. **C and D**. Gradient 1 variance, Gradient 1-4 covariance, and FC differences for two subjects with low or high overall mean atrophy. **E.** Gradient 1-2 covariance and FC differences for two subjects with right parietal or left anterior temporal atrophy. **F.** Gradient 2 variance and FC differences for two subjects with posterior temporal or frontal atrophy.

We then assessed how functional connectivity component F1-F3 scores related to specific across-subject differences in the gradient temporal variance and covariance. Henceforth when describing gradient timeseries, we use the terms “variance” and “amplitude” interchangeably, as well as “covariance” and “phase”. We found that all three function components had strong and specific relationships with the six primary gradients, explaining most of the across-subject FC variance. For F1, the first six gradients explained 82.0% of the functional connectivity variance, with the biggest contributions from Gradient 1 variance (26.4% of F1 variance, t=-17.08; p < 0.001 for all reported terms; Figure 4B**/C**), Gradient 1/4 covariance (15.1% of F1 variance, t=+10.32; Figure 4B**/D**), and Gradient 1/5 covariance (10.8% of F1 variance, t=+10.11). Subjects with higher overall mean atrophy had lower Gradient 1 variance, which because of its unipolar nature (with all regions having positive weights), reflected lower global BOLD signal amplitude. The correlation of Gradient 1 variance and global signal amplitude was r=0.91 (p < 0.001). This explained the weakened global functional connectivity, most extremely in sensory-motor regions with the largest Gradient 1 weights. Subjects with high overall mean atrophy also had stronger Gradient 1-4 covariance than low atrophy subjects. This resulted in more positive functional connectivity between regions with positive weights on Gradient 4 (dorsal attention, cingulo-opercular) and positive weights on Gradient 1 (all regions). In contrast, regions with negative weights on Gradient 4 (default mode) had negative correlation with Gradient 1 and therefore with the whole brain. Thus, atrophy-driven changes in coupling of Gradient 1 with other gradients was mathematically equivalent to specific networks increasing or decreasing their integration with the rest of the brain.

Function components 2 and 3 also had strong relationships with gradient activity. Gradient activity explained 86.2% of the F2 variance, with the strongest influence by Gradient 1 variance (58.3% of variance, t=+31.02), Gradient 1/2 covariance (10.8% of variance, t=+10.82; Figure 4B**/E**), and Gradient 1/6 covariance (5.3% of variance, t=-8.93). F3 had 55.2% of its variance explained by Gradient 1/2 covariance (14.0% of variance, t=-8.20; Figure 4B**/F**), Gradient 2 variance (10.2% of variance, t=-6.97), and Gradient 2/3 covariance (6.8% of variance, t=-5.21). Here, subjects with greater frontal atrophy had lower Gradient 2 variance, reflecting reduced within-network FC for regions on either pole of the gradient, and also less anticorrelation (i.e. stronger FC) between regions at opposite gradient poles. Overall, the majority of the atrophy-associated FC variance was explained by six gradients and their interactions, indicating a low-dimensional basis for the observed FC alterations.

### Gradient phase and amplitude changes reflect hypo and hyperconnectivity patterns

The observation that atrophy and FC were associated with gradient activity prompted a question: can we simulate these altered FC patterns with a generative model and if so, what can that model tell us about how atrophy disrupts brain activity dynamics? We modeled gradient interactions as a system of linear coupled harmonic oscillators (**Methods**), based on our previous work showing that different FC patterns can be generated by spatial gradients interacting via specific coupling parameters ^30^. Eigendecomposition of a coupled oscillator model yields a set of eigenmodes that capture the system dynamics and are useful for analyzing perturbations (Figure 5A). Each eigenmode represents one spatio-temporal component of the system, oscillating at a single fixed frequency. On a given eigenmode, each gradient has a specific phase angle and amplitude. Overall brain activity at each timepoint can then be represented by the summed activity of the gradients across eigenmodes. Here, we computed the gradient coupling parameters for each subject and derived eigenmodes. We validated the accuracy of this modeling approach by simulating gradient timeseries based on subject-specific eigenmodes and comparing simulated to actual FC patterns. We found that each subject’s simulated and actual FC was significantly more similar than to simulated FC from the other subjects (self-actual vs. self-simulated, median r=0.96; self-actual vs. other-simulated, median r=0.72; t=31.46, p < 0.001; **Supplementary Figure 7**). This supported our use of eigenmode analysis for quantifying individual differences in gradient coupling that gave rise to FC differences.

**Figure 5.**
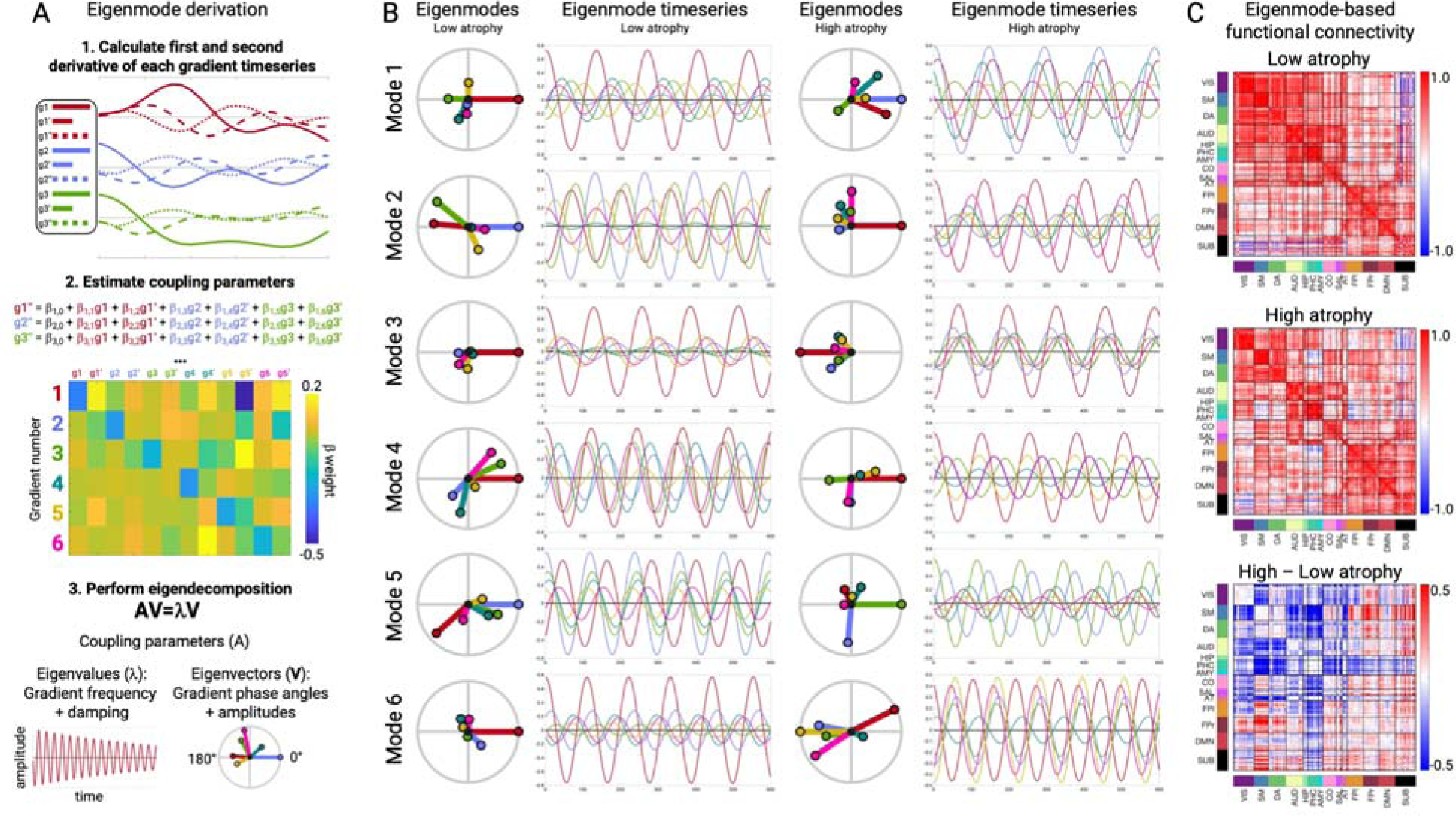
Eigenmode derivation and properties in low and high atrophy subjects. **A.** Procedure for deriving gradient coupling parameters and eigenmodes. **B.** The six eigenmodes for two example subgroups (*n*=5) with the lowest or highest total atrophy. Eigenmodes are ordered from lowest to highest frequency. Circle plots show the phase angle and amplitude of each gradient on each eigenmode. Timeseries plots show the resultant gradient oscillations, occurring at the eigenmode-specific frequency. **C.** Eigenmode-based FC matrices for low and high atrophy subjects and the FC difference matrix.

As an illustration of how the eigenmodes captured different FC patterns, we computed the eigenmodes for two groups of subjects (*n*=5) with the lowest or highest overall mean atrophy as measured by structure-function component 1 (Figure 5B). We used these group-specific eigenmodes to simulate gradient timeseries and compute FC matrices (Figure 5C). The simulated FC differences between low and high atrophy subjects revealed the same pattern of atrophy-associated FC changes represented on SF1 – reduced primary sensory FC, elevated subcortical-cortical FC, and elevated fronto-parietal association FC. This indicated that the eigenmodes contained sufficient information to explain the observed hypo/hyper-FC patterns.

Based on the demonstrated relationship between eigenmodes and atrophy-associated FC patterns, we hypothesized that two eigenmode-derived quantities would capture the atrophy-associated FC alterations: 1) the net amplitude of each gradient across all six eigenmodes and 2) the average phase angle between each pair of gradients across all modes. We statistically evaluated this by measuring the correlation between eigenmode gradient amplitude/angle and gradient variance/covariance. These quantities were strongly related (**Supplementary Figure 8**; corresponding: median absolute r=0.77, p=6.21×10^-65^; non-corresponding: median r=0.07, p=0.19). This demonstrated that atrophy-related shifts in brain-wide FC corresponded to decreasing gradient amplitudes and alterations in the typical phase angle between pairs of gradients.

We further examined the gradient amplitude and phase angle properties for four gradient relationships most strongly associated with the structure-function components. Subjects with lower Gradient 1 variance had either higher overall mean atrophy (SF1) or right parietal atrophy (SF2), which significantly correlated with Gradient 1 amplitude (r=0.66, p < 0.001; Figure 6A). This reduced amplitude resulted in globally reduced functional connectivity strength because of Gradient 1’s unipolarity. Subjects with high overall mean atrophy also had increased Gradient 1-4 covariance, which negatively correlated with Gradient 1-4 phase angle (r=-0.80, p < 0.001; Figure 6B). The phase angle progressively decreased from 93° for subjects in the 20^th^ percentile to 64° for subjects in the 80^th^ percentile. This smaller angle reflected more positive temporal correlation between the Gradient 1 and 4 timeseries, resulting in hyperconnectivity of regions with the same sign on each gradient (+/+ or -/-) and hypoconnectivity of regions with opposite signs (+/-). We interpreted a shift away from 90° as a “collapse”, given that the gradients were identified as temporally orthogonal (i.e. uncorrelated) components in cognitively normal control subjects. For Gradients 1 and 2, there was again a negative correlation between covariance and phase angle (r=-0.80, p < 0.001; Figure 6C). In this case, the collapse away from 90° was in one of two directions, depending on the atrophy pattern. Subjects with posterior temporal atrophy (SF3) or anterior temporal atrophy (SF2) showed an increase to 109° (10^th^ percentile) while subjects with frontal or right parietal atrophy showed a decrease to 57° (90^th^ percentile). This bi-directional collapse resulted in opposite patterns of hypo and hyperconnectivity, resulting from regions with positive or negative weights on Gradient 1 and 2 coming into phase or going out of phase. Finally, Gradient 2 variance significantly correlated with amplitude (r=0.74, p < 0.001; Figure 6D). Subjects with posterior temporal atrophy (SF3) had greater Gradient 2 amplitude than those with frontal atrophy. This reflected more extreme correlated fluctuations for regions with the same sign on Gradient 2 (+/+, -/-) and anticorrelated fluctuations for regions with opposite signs (+/-). Overall, eigenmode analysis revealed that each atrophy-associated hypo/hyperconnectivity pattern was linked to specific gradient amplitude and phase angle alterations.

**Figure 6.**
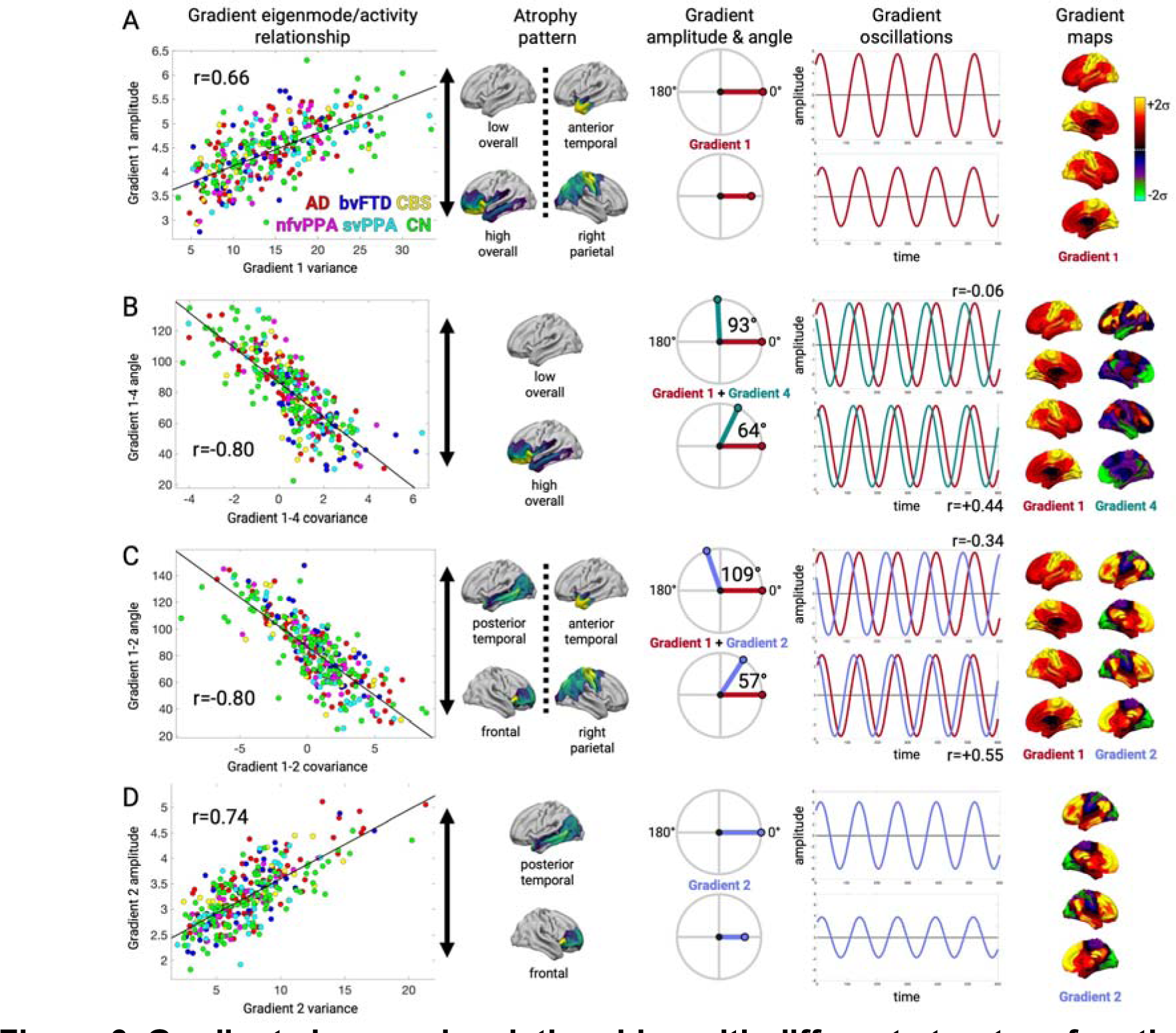
Gradient eigenmode relationships with different structure-function components. **A-D.** Left to right: the correlation between each subject’s gradient variance/covariance and gradient amplitude/angle; the atrophy components associated with higher or lower gradient amplitude/angle; illustration of the gradient amplitude/angle differences and the resultant simulated timeseries; and the gradient spatial maps. **A.** Gradient 1 amplitude associated with atrophy component 1. Gradient amplitudes and timeseries are shown for subjects in the 20^th^ and 80^th^ percentile of atrophy component 1. **B.** Gradient 1-4 angle associated with atrophy component 1. Gradient phase angles and timeseries are shown for subjects in the 20^th^ and 80^th^ percentile of atrophy component 1. R-values show correlation between the simulated gradient timeseries. **C.** Gradient 1-2 angle associated with atrophy components 3 and 2. Gradient phase angles and timeseries are shown for subjects in the 10^th^ and 90^th^ percentile. **D.** Gradient 2 amplitude associated with atrophy component 3. Gradient amplitudes and timeseries are shown for subjects in the 20^th^ and 80^th^ percentile of atrophy component 3.

## Discussion

Here we analyzed structural and task-free functional MRI scans from patients across the AD-FTD spectrum to identify relationships between gray matter atrophy and functional connectivity. We discovered three reproducible structure-function components. The primary component represented a relationship between overall mean atrophy, regardless of spatial location, hypo-connectivity in primary cortical regions, and hyper-connectivity in subcortical and fronto-parietal association cortex regions. The second and third components linked focal syndrome-specific atrophy patterns to peri-lesional hypo-connectivity and distal hyper-connectivity. These structural and functional alterations collectively contributed to impairments in global and domain-specific cognition. Each functional component could be accounted for by variance in six intrinsic activity gradients, suggesting that the disease-related functional alteration patterns are constrained by the brain’s intrinsic functional architecture. Eigenmode analysis of the gradient temporal dynamics revealed reduced amplitude of specific gradients and collapsed phase angles between gradients, offering a possible explanation for the observed patterns of hypo and hyperconnectivity.

Three brain atrophy components explained two-thirds of the variance in atrophy across this diverse set of AD and FTD syndromes that collectively involved nearly all cortical and subcortical regions. The first atrophy component captured mean overall mean atrophy and served as a proxy for disease stage. The second and third components described opposing focal atrophy patterns in the left-predominant temporal pole versus right-predominant dorsal parietal cortex (component 2) and left-predominant temporal-occipital areas versus prefrontal-insula-cingulate areas (component 3). These components stratified patients with different syndromes, separating atrophy patterns along ventral-dorsal, anterior-posterior, and left-right axes. Atrophy subtypes have been well-characterized in different AD and FTD syndromes ^37,38,5,39–43^. We used partial least squares regression to derive atrophy components and continuous scores so that we could link progressive atrophy patterns across stages to corresponding functional connectivity alterations. It may not always be appropriate to represent patients with AD and FTD along such a continuum, as these syndromes are often caused by distinct neuropathological diseases that involve specific cell populations, subcortical nuclei, or cortical layers ^44–47^. Future work may assess structure-function relationships in distinct categorical groups such as those with homogenous underlying pathological substrates.

Distinct functional hypo/hyper-connectivity patterns were associated with each atrophy component. We did not anticipate finding structure-function component 1’s convergent pattern of FC alteration in patients with different syndromes and heterogenous atrophy patterns. However, recent fMRI studies in Parkinson’s disease and Alzheimer’s disease have reported similar patterns of primary sensory connectivity decrease and subcortical and/or association network connectivity enhancements ^33,48,34^, regions that are remote from the primary sites of pathology and neurodegeneration.

There are several ways connectivity can be altered in regions without structural damage. Classical diaschisis involves a focal lesion causing depressed function in a structurally intact target region ^49^, while connectomal diaschisis includes network-wide alterations ^50^. Here we assessed widespread FC alterations by deriving summary scores capturing different FC patterns and then linked these scores to underlying low-dimensional gradient activity levels. We found that the principal FC pattern was based on reduced Gradient 1 variance and increased Gradient 1-4 coupling. A critical question is how atrophy in disparate locations can cause identical convergent alterations in these functional systems. Previous studies have reported widespread FC decreases and increases following focal stroke lesions ^51^, with weak spatial correspondence between structural and functional alterations ^52^. This suggests that focal structural damage can cause non-local disruptions of large-scale network dynamics. Given the evidence that brain activity dynamics occur in a low-dimensional functional state space ^29,20,30^, it may be expected that disparate lesions will converge on a limited range of FC perturbations. In the healthy brain, the optimal set point for each gradient may be temporal orthogonality, maximizing the spatiotemporal segregation of different networks ^53^.

Structural damage may cause specific gradients to collapse away from orthogonality, resulting in hypo- and hypersynchrony between different networks as two sides of the same coin. This may occur in a convergent or divergent fashion, as we found here with SF1 and SF2/3, respectively. The linkage between functional connectivity, activity gradients, and eigenmodes raises a question about whether any of these processes are epiphenomenal. Gradients represent standing waves that combine by superposition to create activity flow patterns described by eigenmodes ^54,55^. Future studies may consider how atrophy perturbs activity flow and whether damaged structural connections play a key role.

The structure-function components explained 25-50% of the variance in cognitive deficits. These deficits clustered into two groups: clinical/global functioning, language, and episodic memory, most strongly influenced by S1, F1, and S3; and executive function, processing speed, language production, and visuospatial processing, driven by S1 and S2. This clustering is consistent with the emerging recognition that diverse brain lesions cause convergent low-dimensional behavioral deficits . The structural components tended to be stronger predictors of cognitive deficits than functional components, which is perhaps unsurprising given that the syndrome diagnostic criteria include structural neuroimaging features. Nonetheless, functional components did explain significant variance in cognitive deficits. Most strikingly, F1 was a significant predictor of CDR-SB over and above the amount of overall mean atrophy (S1 score). F1 may represent a neural substrate for cognitive reserve that can compensate for structural degeneration ^57^. A consistent finding across dementia syndromes is that functional connectivity can compensate for atrophy or molecular pathology to preserve cognitive functioning in FTD and AD ^21,58,59^. Future work may examine whether subjects with worse-than-expected function for their amount of atrophy, as measured by the residuals from the structure-function regression line, may have more reserve and greater potential response to treatment. Our assessment of structure-function across disease stages had several important limitations. We did not consider staggered relationships between biomarker measures ^60^. We did not include patients with mild cognitive impairment or presymptomatic disease and did not assess potential biphasic relationships between atrophy and FC ^61,62^. We also did not attempt to identify structure-function components related with typical aging ^63,64^. Finally, this study only considered task-free fMRI, leaving open how task-engaged brain activity is disrupted by these atrophy patterns.

Function component 1 (F1) correlated with clinical impairment equally well in FTD and AD. F1 has several desirable biomarker properties ^65^ including: 1) surrogacy with CDR, the standard primary endpoint in dementia clinical trials ^66^; 2) longitudinal within-subject correlation with clinical worsening and progressive atrophy; and 3) reproducible relationships with brain structure and clinical severity scores across 37 different sites combining the UCSF and ADNI datasets. While molecular and anatomical neuroimaging biomarkers are more widely applied in late-stage dementia trials ^67^, fMRI biomarkers have the potential to measure cognition-supporting brain activity with high anatomical precision prior to widespread neurodegeneration. In this study, structural and functional components explained independent variance in CDR. This suggests patients might benefit from treatments that slow neurodegeneration, restore function, or both, and monitoring both structural and functional biomarkers could add value to clinical trials. While the current observational study in a diverse cohort was well suited for biomarker discovery, a key next step is analytical and clinical biomarker validation.

This effort should focus on specified contexts of use, including as a predictive biomarker for identifying individuals more likely to respond to treatment, or as a monitoring biomarker for detecting treatment response ^68^. This will ideally include an fMRI acquisition protocol that optimizes within-subject reliability ^69,70^. It will also be prudent to consider the effect of symptomatic therapies on activity imbalance, given that activity gradients reflect neurotransmitter receptor distributions ^29,71,30^ and may be modulated by acetylcholinesterase inhibitors or selective serotonin reuptake inhibitors commonly used in dementia treatment.

## Methods

### Subject selection

Patients with dementia and cognitively normal control subjects were recruited through ongoing studies at the University of California San Francisco (UCSF) Memory and Aging Center. All subjects or their surrogates provided informed consent according to the Declaration of Helsinki and the procedures were approved by the UCSF Institutional Review Board. All subjects underwent a clinical history, physical examination, neuroimaging, and neuropsychological assessment within 90 days of scanning. Cognitively normal control subjects were recruited from the Hillblom Healthy Aging Study with ages between 45-85, minor or no memory problems, a clinical dementia rating score of 0, and no diagnosis of a neurodegenerative disease or other major health condition. Subjects were excluded if they had significant history of other neurological diseases or structural brain abnormalities inconsistent with their primary clinical syndrome. Subjects were included whether or not they took medication for symptoms of Alzheimer’s disease or frontotemporal dementia.

A large initial control dataset (n=568) was subsequently divided for multiple purposes. The initial patient group consisted of patients (n=309) who received a primary clinical diagnosis of either Alzheimer’s disease ^72^, behavioral variant frontotemporal dementia ^73^, semantic variant or nonfluent variant primary progressive aphasia ^74^, or corticobasal syndrome ^75^. All diagnoses were made within 90 days of the patients’ MRI scan. We assigned each patient to a high, intermediate, or low confidence diagnosis group. High confidence subjects had a single clinical diagnosis of one of the five syndromes of interest at one or more clinical visits. Intermediate confidence subjects had a best estimate clinical diagnosis of the syndrome of interest, but additionally either: 1) an alternative possible diagnosis including AD, bvFTD, CBS, nfvPPA, svPPA, progressive supranuclear palsy (PSP), amyotrophic lateral sclerosis (ALS), posterior cortical atrophy (PCA), or logopenic variant PPA (lvPPA) or 2) a best estimate clinical diagnosis that was stable for multiple visits before shifting away from the syndrome of interest in a later visit. These subjects were assigned to the intermediate confidence group if they had three or more clinic visits with a stable primary diagnosis of the syndrome of interest. Low confidence subjects had multiple diagnoses as best estimates, including the syndrome of interest, or a diagnosis for the syndrome of interest that shifted away at the next visit. Our primary analysis focused on 221 high and intermediate confidence patients with dementia, excluding low confidence patients and MRI quality control failures (see below). 100 cognitively normal subjects were selected who passed image quality control and were matched to the overall patient group for age, sex, MRI scanner distribution, and fMRI head motion. The number of subjects with each diagnosis are shown in **Table 1**. The self-reported race/ethnicity for the 321 subjects included 25 Asian, 5 Black, 5 Hispanic, 263 White, and 23 unreported.

### Neuroimaging acquisition

All subjects were scanned at the UCSF Neuroscience Imaging Center, on either Siemens Trio or Siemens Prisma Fit 3T MRI scanners. Subjects were scanned between one and eight times over the course of their clinic visits. Our main analysis focused on the baseline scan for each subject. The number of patients with each diagnosis scanned on either the Trio or Prisma are shown in **Table 1**. Subjects received T1-weighted magnetization-prepared rapid gradient echo structural MRI (MPRAGE) scans with similar acquisition parameters on the Trio or Prisma: acquisition time: 8:53; sagittal slice orientation; thickness: 1.0 mm; field of view: 160×240×256 mm; isotropic voxel resolution: 1mm^3^; TR: 2300 ms; TE: 2.98 ms for Trio, 2.9 ms for Prisma; TI: 900 ms, flip angle: 9°.

Task-free fMRI scans were run using a T2*-weighted echoplanar scan with subjects instructed to remain awake with their eyes closed. The parameters on the Trio were: acquisition time: 8:06; axial orientation with interleaved ordering; field of view: 230×230×129 mm; matrix size: 92×92, effective voxel resolution: 2.5×2.5×3.0 mm; TR: 2000 ms, for a total of 240 volumes; TE: 27 ms. For the Prisma, the fMRI parameters were: acquisition time: 8:05; axial orientation with interleaved multi-slice mode and multiband acceleration=6; field of view: 211×211×145 mm; matrix size: 92×92, effective voxel resolution: 2.2×2.2×2.2 mm; TR: 850 ms, for a total of 560 volumes; TE: 33 ms.

### Structural image processing

MPRAGE scans were visually assessed by trained technicians and scans with excessive motion artifact (ringing or blurring) were excluded. MPRAGE scans for all time points for a given subject that passed visual inspection were registered using the serial longitudinal registration in SPM12 ^76^. Default parameters were used for warping regularization and bias regularization. Jacobian determinant and divergence maps were produced that represent the amount of longitudinal brain contraction and expansion. We then applied unified normalization/ segmentation to register the midpoint average T1 images to the MNI152NLin6Asym standard space ^77^ with light regularization, a 60 mm bias FWHM cutoff, and Gaussians per tissue type of [2,2,2,3,4,2]. The gray matter tissue segmentation for the midpoint average was multiplied by the deformation fields for each time point to obtain time point-specific gray matter maps. These images were then warped to standard space using the deformation fields from the unified normalization/segmentation procedure. The resulting normalized gray matter maps were smoothed with an 8 mm FWHM Gaussian kernel.

We derived voxelwise gray matter tissue probability maps from an independent set of cognitively normal control subjects (n=397) using the same structural image processing methods. These subjects had the following characteristics: mean age=69.3±8.8; 239 female/158 male; 345 right-handed/43 left-handed; 1.5T/3T Trio/3T Prisma/4T=58/144/140/52). We ran multiple regression for each voxel to estimate gray matter volume as a function of age, sex, handedness, total intracranial volume, and MRI scanner identity. For the 321 subjects in the primary analysis, we entered their demographic values into this regression model to estimate their gray matter volume in each voxel. The voxel W-score was calculated as the difference between actual gray matter volume and the estimated gray matter volume, divided by the standard deviation of the model fit in the reference control sample (LaJoie et al., 2012). W-scores were used as the measurement of gray matter atrophy throughout the study.

### Functional image processing

Functional MRI scans were processed using fMRIPrep (Esteban et al., 2019) (RRID:SCR_016216). For anatomical image processing, the MPRAGE images were corrected for intensity non-uniformity with N4BiasFieldCorrection in ANTs (Avants et al., 2008) (RRID:SCR_004757), and used as the T1-weighted (T1w) reference throughout the workflow. The T1w reference was skull-stripped with a Nipype ^79^ (RRID:SCR_002502) implementation of the antsBrainExtraction.sh workflow using OASIS30ANTs as target template. Brain tissue segmentation of cerebrospinal fluid (CSF), white-matter (WM) and gray-matter (GM) was performed on the brain-extracted T1w using FSL fast (https://fsl.fmrib.ox.ac.uk/fsl/fslwiki; RRID:SCR_002823). Volume-based spatial normalization to the MNI152NLin6Asym standard space was performed through nonlinear registration with antsRegistration, using brain-extracted versions of both T1w reference and the T1w template.

For functional image processing, the first five volumes were removed to allow for scanner stabilization. A reference volume and its skull-stripped version were generated by fMRIPrep. The BOLD reference was then co-registered to the T1w reference using FSL flirt with 6-degrees-of-freedom affine registration. Co-registration was configured with nine degrees of freedom to account for distortions remaining in the BOLD reference. Head-motion parameters with respect to the BOLD reference (transformation matrices, and six corresponding rotation and translation parameters) were estimated using FSL mcflirt and were used to compute the framewise displacement (FD). BOLD runs were slice-time corrected using AFNI 3dTshift (https://afni.nimh.nih.gov/; RRID:SCR_005927). The BOLD images were realigned from native to MNI152NLin6Asym standard space using antsApplyTransforms, configured with Lanczos interpolation, with a single interpolation step by composing transformations for head-motion and co-registrations to anatomical and output spaces. Images were spatially smoothed with a 6mm FWHM (full-width half-maximum) kernel using FSL susan. Confounding CSF and WM timeseries were calculated based on the preprocessed BOLD images, deriving average signals using the subject-specific anatomically derived tissue masks after erosion. The confound timeseries for head motion estimates, CSF, and WM were expanded to include the temporal derivatives and quadratic terms ^80^. Bandpass filtering in the frequency range 0.008-0.08 Hz was performed on the confound timeseries and BOLD images using fslmaths and AFNI 3dBandpass respectively. We did not perform global signal regression and instead assessed global signal variance as a disease-relevant variable of interest. The global signal was computed as the mean BOLD signal across the 246 regions (see below) at each timepoint. Confound timeseries were then regressed out of the BOLD images using fslglm. Scans were standardized voxelwise to have mean=0 and standard deviation=1 across time. Subjects with greater than 0.55 mm mean FD were excluded from subsequent analysis ^81^. From the pool of all fMRI scans for all available control and patient subjects (n=1591), this resulted in the exclusion of 194 scans (12.2%). We performed a subsequent data-driven denoising procedure using PCA to remove scans with implausible functional connectivity patterns likely due to noise from the scanner hardware (field instabilities) and the subject (head motion, heartbeat, respiration). PCA has previously been applied for outlier detection in fMRI data ^82^. Here, all subject’s functional connectivity matrices (see below) were flattened and combined into one single matrix ([1591 scans x 30135 edges]). PCA was run on this matrix and subjects with an outlying score on the first component (> 1 standard deviation above the mean) were flagged. This excluded 288 scans and left an available pool of 1108 scans.

### Scanner harmonization

Structural and functional MRI data from the two different MRI scanners was harmonized using ComBat ^83^. Gray matter mean W-score values were estimated for 246 regions of interest (210 cortical, 36 subcortical including the caudate, putatmen, globus pallidus, and thalamus) from the Brainnetome atlas ^84^. A design matrix was constructed including MRI scanner as the main batch variable and covariates for patient/control status, age, and sex. ComBat was then used to harmonize the W-score values for each region. For functional MRI, mean BOLD timeseries for each scan were obtained for each region and entered into a temporal PCA (see next section). Using the 246 temporal component timeseries, we computed the covariance matrix. The upper triangle from each [246 x 246] covariance matrix was extracted, including the diagonal, flattened into a [30381 x 1] vector, and stacked for all subjects. The [321 x 30381] matrix was run through ComBat with the same design matrix. The harmonized covariance values were then used to derive harmonized FC matrices.

### Gradient derivation and functional connectivity analysis

We studied the low-dimensional basis of functional connectivity patterns by performing dimensionality reduction on fMRI BOLD timeseries data. Specifically, we derived activity gradient spatial maps and temporal activity timeseries based on methods described in our previous work ^30^. Here we obtained an independent cohort of cognitively normal subjects (n=321) that were age, sex, and scanner-matched to our primary cohort of 221 patients and 100 cognitively normal subjects. The 246 regional mean BOLD timeseries for the independent cohort subjects were temporally concatenated into a [122795 x 246] matrix and PCA was performed. This yielded a brain activity latent space in which brain region component loadings (eigenvectors scaled by their corresponding eigenvalues) represented each region’s weight on that component. We refer to the spatial maps for each component as gradients and the component scores as the gradient timeseries. Our main analysis focused on the first six dimensions with additional consideration of 12 dimensions. The fMRI ROI timeseries matrix for the primary cohort ([119915 x 246]) was projected into this latent space to obtain the gradient timeseries. For each subject, we computed the [246 x 246] gradient covariance matrix. We used these matrices to derive functional connectivity matrices by obtaining each region’s variance (summing on-diagonal values across all components), each region pair’s covariance (summing off-diagonal values across all component pairs), and using these quantities to compute the region pairwise Pearson correlation coefficients. An independent PCA was run on the primary cohort [119915 x 246] ROI timeseries matrix to validate the spatial gradient pattern reliability.

### Brain structure-function statistical analysis

The relationship between atrophy and brain-wide functional connectivity was assessed in two stages. In the first stage, we vectorized each subject’s FC matrix upper triangle into a [30135 x 1] vector and stacked these to obtain the ([321 subject x 30135 edge] group FC matrix. We performed partial least squares regression on the brain atrophy W-scores ([321 subjects x 246 regions]) and the functional connectivity ([321 subject x 30135 edges]). PLSR is asymmetric, designed to decompose the ‘X’ variable into components that maximally covary with ‘Y’ variable ^85^. We chose to use atrophy as the ‘X’, i.e. the “grounding” variable, based on two factors: 1) our stronger hypothesis about the spatial patterns of the atrophy components, and 2) lower variability in structural MRI than functional MRI. We ran PLSR to derive five components and used split-half analysis to determine which components had sufficient reliability. Here we focused on the structural component reliability and describe our procedure for functional component reliability in the next section. We generated 1000 random splits of the 321 subjects into two equally sized groups, each time balancing the number of subjects with each clinical syndrome. We ran 1000 trials of independent PLSR on the first and second halves of the subjects. The structure component loadings were compared for each of the 1000 trials using correlation. Analysis revealed that the first three components had acceptable reliability. We obtained the atrophy and FC scores for each subject for these three components and computed the structure-function correlation from these scores to measure the strength of each independent structure-function relationship.

We validated the reliability of the structure-function relationships by performing ridge regression with cross-validation. For this analysis, we could not use atrophy component scores from our PLSR model because these were derived based on their covariance with FC and would contaminate cross-validation. Instead, we derived atrophy component scores using PCA on the atrophy data alone. The spatial patterns of the first three PCA-derived atrophy components were essentially identical to the first three PLSR atrophy components (Component 1, r=0.999; Component 2, r=0.98, Component 3, r=0.97). Ridge regression was run using scikit-learn ^86^ (https://scikit-learn.org/stable/modules/generated/sklearn.linear_model.Ridge.html). We estimated models of the relationship between each atrophy component (one score per subject per component) and brain-wide FC weights (30135 features per subject). Stratified cross-validation was run to measure model out-of-sample accuracy with four folds with 240 train subjects and 81 test subjects per fold, balanced for the number of subjects with each syndrome. We ran 1000 trials with an empirically selected alpha value of 1000, which produced optimal cross-validation accuracy. Ridge coefficients for each component were averaged across 1000 trials and 4 folds to obtain the average coefficients. Individual subject functional scores for each component were derived by taking each subject’s score from their left-out fold for each trial and averaging across the 1000 trials. Deriving these out-of-sample ridge FC scores also served to decorrelate structure and function scores sufficiently to use them as independent predictors of neuropsychological scores.

We visualized structure-function relationships for each component as matrices of PLSR FC edge weights. The 246 regions were grouped into 14 previously defined functional connectivity modules ^87^, based on a modular partitioning of a group-averaged task-free functional connectivity matrix from 75 healthy older control subjects. These modules included in this partition are: visual, sensory-motor, dorsal attention network, auditory, hippocampal, parahippocampal, amygdala, cingulo-opercular, salience, anterior temporal, left fronto-parietal, default mode network, right fronto-parietal, and subcortical. We created spatial maps summarizing the most prominent FC patterns for each component by computing region-wise sums of the PLSR FC edge weight matrices.

Syndrome-associated atrophy and functional connectivity patterns were assessed by identifying the subset of patients expressing the typical pattern. We determined this by performing linear discriminant analysis on structure component 1-3 scores with syndrome label as the response variable. This resulted in 51/82 AD patients, 25/41 bvFTD, 10/27 CBS, 9/34 nfvPPA, 32/37 svPPA, and 76/100 CN. The mean atrophy map and FC matrix were computed for each group and (syndrome – cognitively normal) alterations were derived. The reconstructed FC matrix for each group was computed as the outer product of function component 1-3 scores and the corresponding loadings. Correlations were measured between the typical mean actual FC matrix for each syndrome and the reconstructed FC matrix.

### Dynamical systems modeling

We used dynamical systems modelling to analyze gradient dynamic activity in different groups of subjects ^30^. All these analyses used the first six gradients. For each gradient timeseries, the first and second derivatives were calculated by finite differencing using the ‘gradient’ function in MATLAB. We then ran linear regression for each gradient to estimate its second derivative timeseries (*G’’*) as a function of all six gradients’ timeseries (*G*) and first derivatives (*G’*) along with an intercept. The parameter estimates (coupling parameters) for the 13 terms from each regression were then used to define a system of six coupled second-order ordinary differential equations:

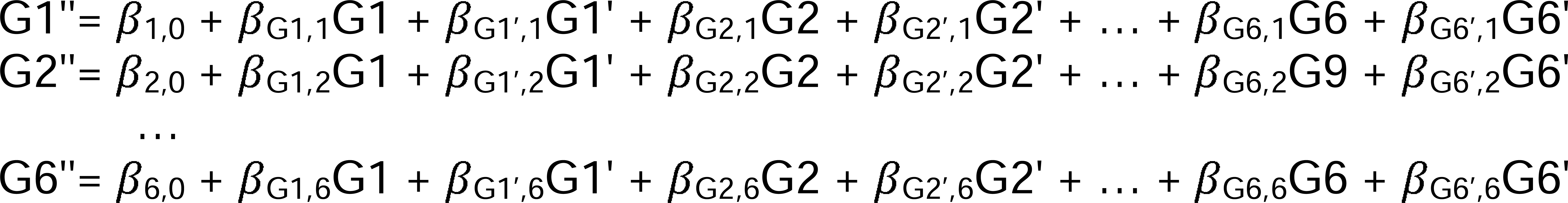

These equations modeled gradient interactions as a system of linear coupled harmonic oscillators with damping ^88^.

Eigendecomposition was used to analyze the governing dynamics of the harmonic oscillator system. We transformed each second order equation to a set of two first order equations using substitution. This linear system can be represented as:

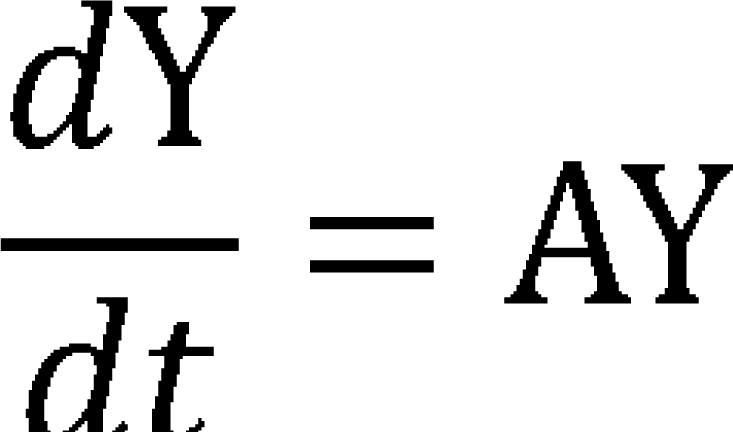

where Y is the gradient timeseries and A is the coupling parameter matrix. Eigendecomposition of the coupling parameter matrix results in:

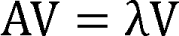

where λ and V are the eigenvalues and eigenvectors for the *m* eigenmodes. Each eigenmode describes a signal oscillating at a specific frequency. Eigendecomposition of a harmonic oscillator system with damping typically yields complex eigenvalues, with the real and imaginary parts (α+*i*β) describing the damping (exponential growth or decay) and frequency of each eigenmode, respectively. The solution to the differential equation with complex eigenvalues is:

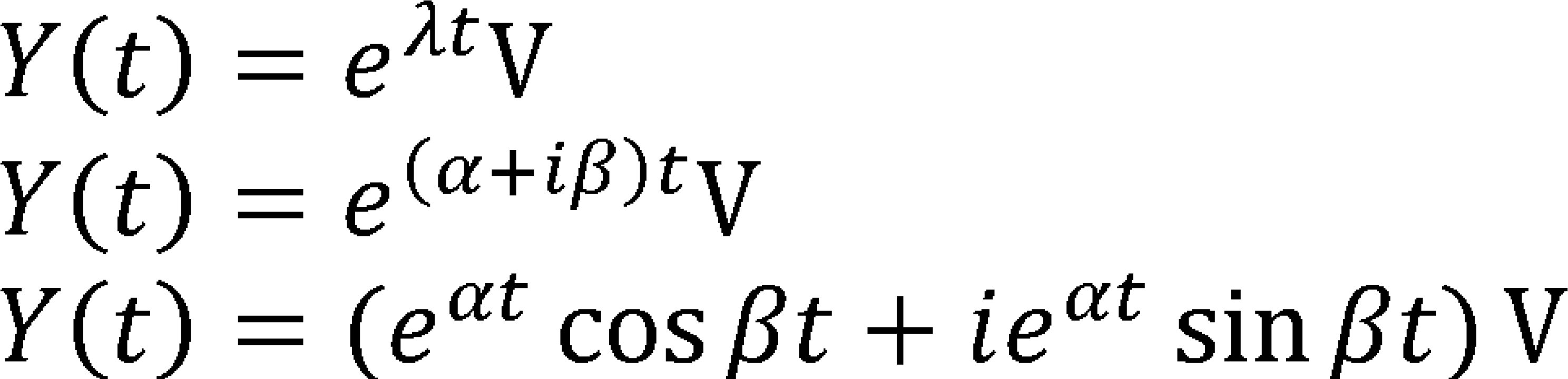

Positive or negative values of α represent the damping of each eigenmode over time.

β is the angular frequency (number of cycles per time unit) and is converted to hertz by β/2π/TR. The eigenvector components are also complex, with the real and imaginary parts describing the amplitude and phase angle of each gradient on that eigenmode. When solving the equation for a given set of initial conditions, the real and imaginary parts of each eigenmode are each scaled by constants *k* to satisfy the initial conditions, e.g.:

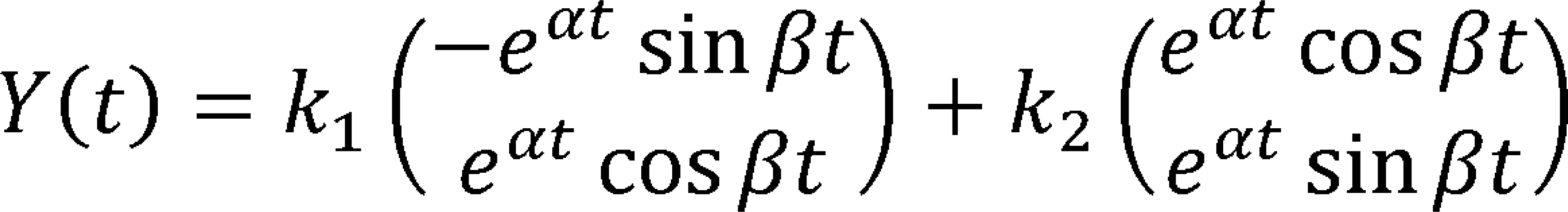

For a given gradient *G*, the overall solution for its timeseries on an eigenmode is:

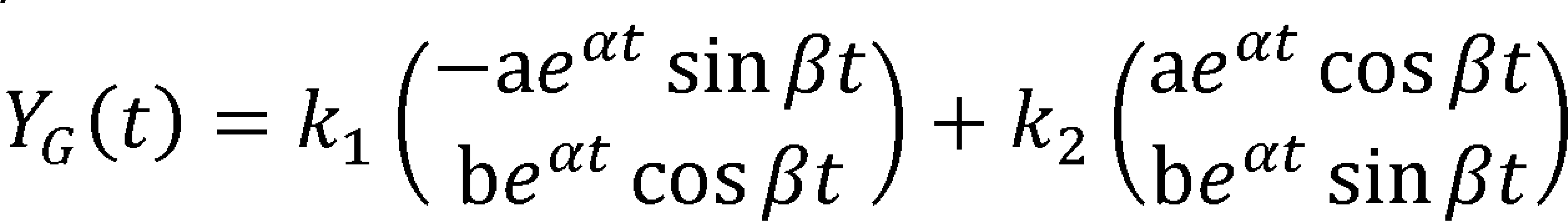

where a and b are the real and imaginary parts of the gradient’s eigenvector component on eigenmode *m*.

The coupling parameter matrix was computed for each subject, from which eigenmodes were derived and gradient timeseries were simulated. These simulations were run using the gradient timeseries and first derivatives for each timepoint as the initial condition and running for the number of timepoints in that subject’s scan (235 or 555 for Trio or Prisma). The [235/555 x 6] gradient timeseries were matrix multiplied by the [6 x 246] region gradient weights to obtain [246 x 235/555] region timeseries, from which [246 x 246] FC matrices were computed. These 235/555 matrices were averaged to produce the subject’s simulated FC matrix. For the actual data, the FC matrices were derived from the six gradients’ timeseries. Simulated and real FC matrices for each condition were statistically compared using Pearson correlation on the matrix upper triangle edge weights.

We performed an illustrative eigenmode analysis on subjects with the lowest and highest function component 1 (F1) scores, associated with the lowest or highest overall mean atrophy. We sorted subjects based on F1 scores and grouped the five subjects with the lowest or highest scores, limited to only subjects with Siemens Trio scans.

Gradient timeseries for the five subjects were concatenated into a [1175 x 6] matrix, from which we computed coupling parameters and derived the eigenmodes. The gradient timeseries were simulated for 1175 timepoints, of which the first 600 timepoints are displayed in Figure 5. The region timeseries and FC matrices were computed and compared for both subject groups.

The eigenmodes for each subject were used to measure two across-eigenmode quantities: 1) each gradient’s cumulative amplitude 2) each gradient pair’s cumulative phase angle. A gradient’s cumulative amplitude was computed as:

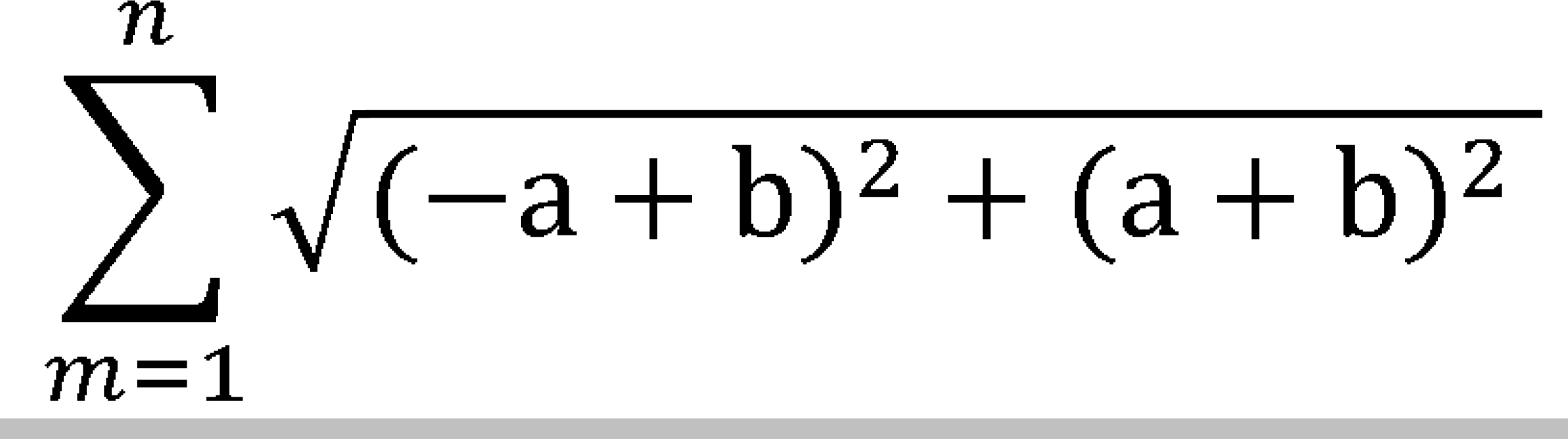

The phase angle difference between a pair of gradients on a given mode was calculated by subtracting the angles for a_1_+*i*b_1_ and a_2_+*i*b_2_. The cumulative phase angle difference between a pair of gradients across eigenmodes was then measured as the circular average of angle differences using the ’circ_mean’ function in the ’circstat’ MATLAB package ^89^, weighted by the product of the gradient amplitudes on the respective eigenmodes. Gradient amplitudes and gradient-pair angles (21 measurements per subject) were then correlated across subjects with gradient variance and covariances (21 measurements per subject), resulting in a [21×21] correlation matrix.

### Neuropsychological testing

Each subject completed a neuropsychological battery. This included assessments for global cognition and function with the Clinical Dementia Rating CDR®+NACC-FTLD ^36^ and the Mini-mental status exam (MMSE) ^90^. The battery also tested the following domains: episodic memory with the short form California Verbal Learning Test (CVLT) ^91^ and the Benson figure delayed recall from the uniform data set ^92^; working memory with the forward and backward digit span length tests; processing speed with the trail making test (Trails A) and Stroop naming tests ^93^; executive function with the Stroop interference test, Delis–Kaplan Executive Function System (DKEFS) Design Fluency ^94^, and the modified trail making test (Trails B); visuospatial processing with the Benson figure copy test ^95^ and Number Location subtest of the Visual Object and Space Perception battery (VOSP) ^96^; and speech and language with the Boston Naming Test (BNT) ^97^, the Peabody Picture Vocabulary Test (PPVT) ^98^, the animal and vegetable naming tests, the D-letter naming test, syntax comprehension ^99^, and verbal articulatory agility ^100^.

### Brain-behavior statistical analysis

Brain-behavior relationships were estimated with generalized additive models using the ‘mgcv’ package ^101^ in R (https://www.r-project.org/). We selected only tests where 120 or more subjects had scores. For each test, we set up a model with predictors of atrophy scores for the first three atrophy components and functional scores associated with the first three atrophy components (from ridge regression). Each of these terms was restricted to have a maximal non-linear basis of 3 by setting k=3 in the model formula. We did not include explicit structure-by-function interaction terms because non-linear terms estimated by a GAM implicitly capture differences in slope across the range of each variable. Each model included linear covariates for age, sex, years of education, scanner type (Trio vs Prisma), and fMRI head motion (mean framewise displacement). Models were fit for 24 cognitive tests and predictors were deemed significant when their F-statistic associated p-values survived global FDR correction (24 tests x 6 brain-based predictors = 148 terms) with q=0.05.

A targeted longitudinal model was used to estimate the relationship between F1, S1, and CDR. 53 subjects (47 patients, 6 cognitively normal subjects) had longitudinal data (mean visit interval=1.1±0.5 years, range=0.4-2.6 years) that passed all quality control tests. S1 and F1 scores were measured for follow-up scans using the linear weights from atrophy PLSR component S1 for structure and from ridge regression for function. We specified a mixed effects model using ‘mgcv’ with CDR-SB score from each scan-associated visit (106 total) as the outcome variable. The model estimated both between-subject and within-subject variation as in ^62^ by including S1 subject mean (averaged across timepoints; non-linear basis k=3 to match the cross-sectional model), S1 longitudinal change (difference from the subject’s mean, varying within-subject across timepoints; k=1 to limit model degrees of freedom), F1 mean (k=3), F1 longitudinal change (k=1), age, sex, years of education, and random intercepts for each subject. In a complementary model, F1 was estimated as a function of S1 baseline (k=3), S1 change (k=3), age, sex, years of education, visit interval, and random intercepts for each subject.

### Replication analysis

A replication dataset was compiled from subjects in the ADNI3 study ^102^, obtained from the Alzheimer’s Disease Neuroimaging Initiative (ADNI) database (https://adni.loni.usc.edu/). The ADNI was launched in 2003 as a public-private partnership, led by Principal Investigator Michael W. Weiner, MD. The primary goal of ADNI has been to test whether serial magnetic resonance imaging (MRI), positron emission tomography (PET), other biological markers, and clinical and neuropsychological assessment can be combined to measure the progression of mild cognitive impairment (MCI) and early Alzheimer’s disease (AD). For up-to-date information, see www.adni-info.org. We included subjects with a diagnosis of cognitively normal or Alzheimer’s disease dementia who received structural MRI and resting state functional MRI scans. A total of 966 visits from 573 subjects (CN n=500, AD n=73) at 68 sites were obtained. We included the subset of scans that were from sites with 10 or more subjects, and that satisfied fMRI motion criteria (mean framewise displacement <= 0.55 mm), resulting in 821 scans. These subjects had the following characteristics: diagnosis, CN n=421, AD n=56; mean age, CN=72.9±8.1 years, AD=75.6±8.1 years; 277 female/200 male; 437 right-handed/40 left-handed; mean interscan interval=1.75±0.7 years). Scans were run on 3T scanners including Siemens (Prisma, n=441; Biograph, n=5; Skyra, n=12; Trio, n=48; Verio, n=91), Philips (Medicare Ingenia, n=54; Achieva, n=78), or GE (Medical Systems Discovery, n=209; Signa n=23). The typical sMRI acquisition parameters were acquisition time: 6:20; sagittal slice orientation; thickness: 1.0 mm; field of view: 208×240×256 mm; isotropic voxel resolution: 1mm^3^; TR: 2300 ms; TE: 3 ms; TI: 900 ms. The typical single-band tf-fMRI acquisition parameters were acquisition time: 10:00; axial orientation with interleaved ordering; field of view: 220×220×163 mm; matrix size: 92×92, effective voxel resolution: 2.2×2.2×2.2 mm; TR: 3000 ms, for a total of 560 volumes; TE: 30 ms; with instructions to remain awake with eyes open. The typical multi-band tf-fMRI acquisition parameters were acquisition time: 10:00; axial orientation with interleaved multi-slice mode and multiband acceleration=8; field of view: 220×220×160 mm; matrix size: 92×92, effective voxel resolution: 2.5×2.5×2.5 mm; TR: 670 ms; TE: 30 ms. We included n=189 multiband scans (TR=0.67/0.79ms, only from Siemens Prisma/Prisma Fit/Syra scanners) and n=777 single-band scans (TR=3/3.15s). CDR-SB scores were available for 774/821 scans. Images were processed with the same pipelines used for the main dataset. Atrophy W-maps and [246 x 1] region atrophy vectors were derived for each structural scan using the same W-score model with the default intercept. [246 x 246] FC matrices were obtained for each functional scan. We harmonized atrophy and FC data across sites by running ComBat on the atrophy/FC values for all scans, controlling for patient/control status, age, and sex. We then computed atrophy component scores for components 1-3 using the atrophy PCA loadings. Functional connectivity component scores were computed for components 1-3 using the average ridge regression coefficients. Structure-function relationships were estimated using a mixed effects linear model with functional component score as the response variable, fixed effects for the corresponding structural component score, mean age, sex, years of education, and framewise displacement, and random intercepts for subject and site. CDR scores were estimated using a mixed effects GAM with CDR-SB as the response variable, non-linear fixed effects for S1-S3 and F1-F3 (all with a non-linear basis k=3), linear fixed effects for mean age, sex, years of education, and framewise displacement, and random intercepts for site.

## Acknowledgements

This work was supported by NIH grants K01AG055698 (J.A.B.), K99-AG065457 (L.P.), R00AG065457 (L.P.) P50AG023501, P01AG019724, the Tau Consortium, the Bluefield Project to Cure FTD, and the Larry L. Hillblom Network Grant for the Prevention of Age-Associated Cognitive Decline. We are grateful for the participation of the patients and their caregivers who made this work possible. Replication data collection and sharing for this project were funded by the Alzheimer’s Disease Neuroimaging Initiative (ADNI) (National Institutes of Health Grant U01 AG024904) and DOD ADNI (Department of Defense award number W81XWH-12-2-0012). ADNI is funded by the National Institute on Aging, the National Institute of Biomedical Imaging and Bioengineering, and through generous contributions from the following: AbbVie, Alzheimer’s Association; Alzheimer’s Drug Discovery Foundation; Araclon Biotech; BioClinica, Inc.; Biogen; Bristol-Myers Squibb Company; CereSpir, Inc.; Cogstate; Eisai Inc.; Elan Pharmaceuticals, Inc.; Eli Lilly and Company; EuroImmun; F. Hoffmann-La Roche Ltd and its affiliated company Genentech, Inc.; Fujirebio; GE Healthcare; IXICO Ltd.; Janssen Alzheimer Immunotherapy Research & Development, LLC.; Johnson & Johnson Pharmaceutical Research & Development LLC.; Lumosity; Lundbeck; Merck & Co., Inc.; Meso Scale Diagnostics, LLC.; NeuroRx Research; Neurotrack Technologies; Novartis Pharmaceuticals Corporation; Pfizer Inc.; Piramal Imaging; Servier; Takeda Pharmaceutical Company; and Transition Therapeutics. The Canadian Institutes of Health Research is providing funds to support ADNI clinical sites in Canada. Private sector contributions are facilitated by the Foundation for the National Institutes of Health (www.fnih.org). The grantee organization is the Northern California Institute for Research and Education, and the study is coordinated by the Alzheimer’s Therapeutic Research Institute at the University of Southern California. ADNI data are disseminated by the Laboratory for Neuro Imaging at the University of Southern California.

---

**Supplementary Figure 1.**
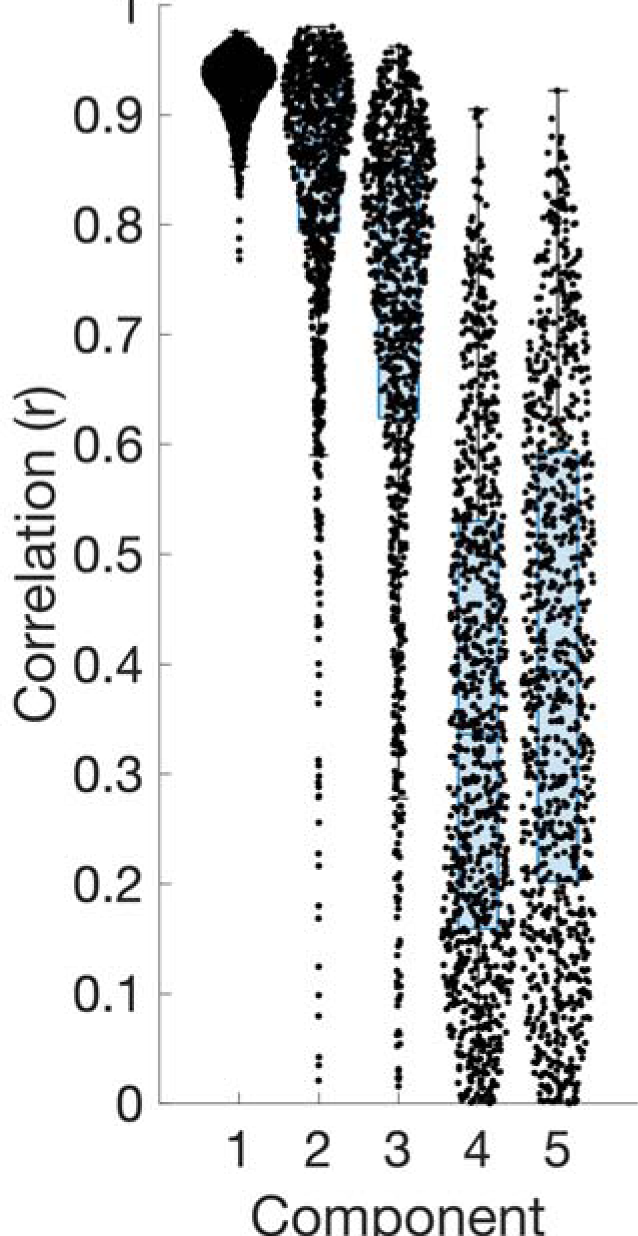
Partial least squares regression structure component reliability using split-half analysis. Supplementary Results 1. 1000 trials of independent PLSR were run on the first and second halves of the subjects, randomly split with balanced syndrome classes for each trial. The structure component loading vectors were correlated between split halves for the 1000 trials for the first five PLSR components. The median correlations were S1: r=0.93±0.03, S2: r=0.88±0.14, S3: r=0.77±0.20, S4: r=0.34±0.23, S5: r=0.39±0.24. The most substantial drop in reliability was between components 3 and 4.

**Supplementary Figure 2.**
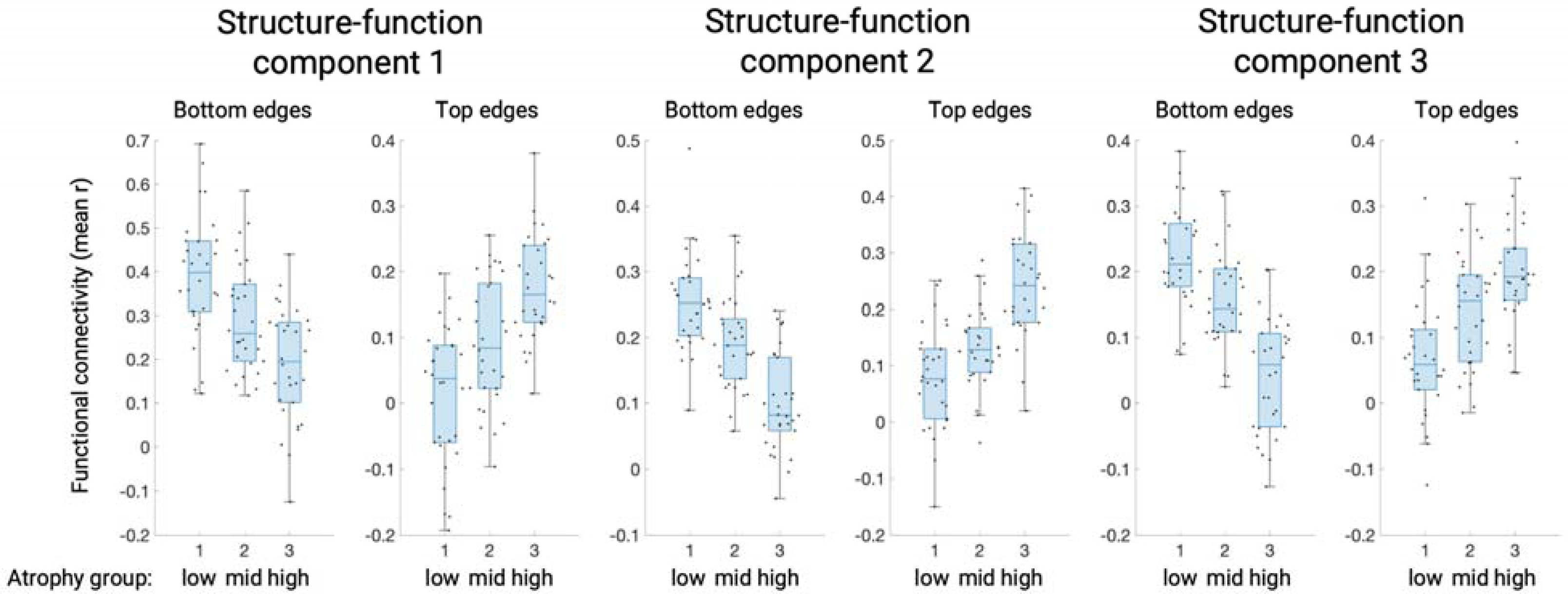
FC edge weights for the bottom/top 1% of edges on each function component. Mean FC edge weights are shown for each component for groups of 30 subjects with the lowest/intermediate/highest atrophy scores. All boxplots show the median, lower and upper quartile range, and the non-outlier minimum/maximum. Supplementary Results 2. For the first three structure-function components, subjects were sorted based on their structural score for that component and binned into groups of 30. For a given component, FC edges were sorted based on their PLSR weight. The bottom/top 1% of edges (300 edges) were kept and the mean FC weight for these two sets were computed for each subject. These mean edge weights were statistically compared for the groups of 30 subjects with low/middle/high atrophy on that component, both for the bottom and top FC edges. The edge weights were always statistically significant between low and high subjects (all p < 0.001, see table) and were significant at a less stringent threshold (p < 0.01) in all other tests. This indicated that the partial FC variance captured by the FC scores was substantial enough to capture significant differences in overall FC edge weights between groups of subjects.

**Supplementary Figure 3.**
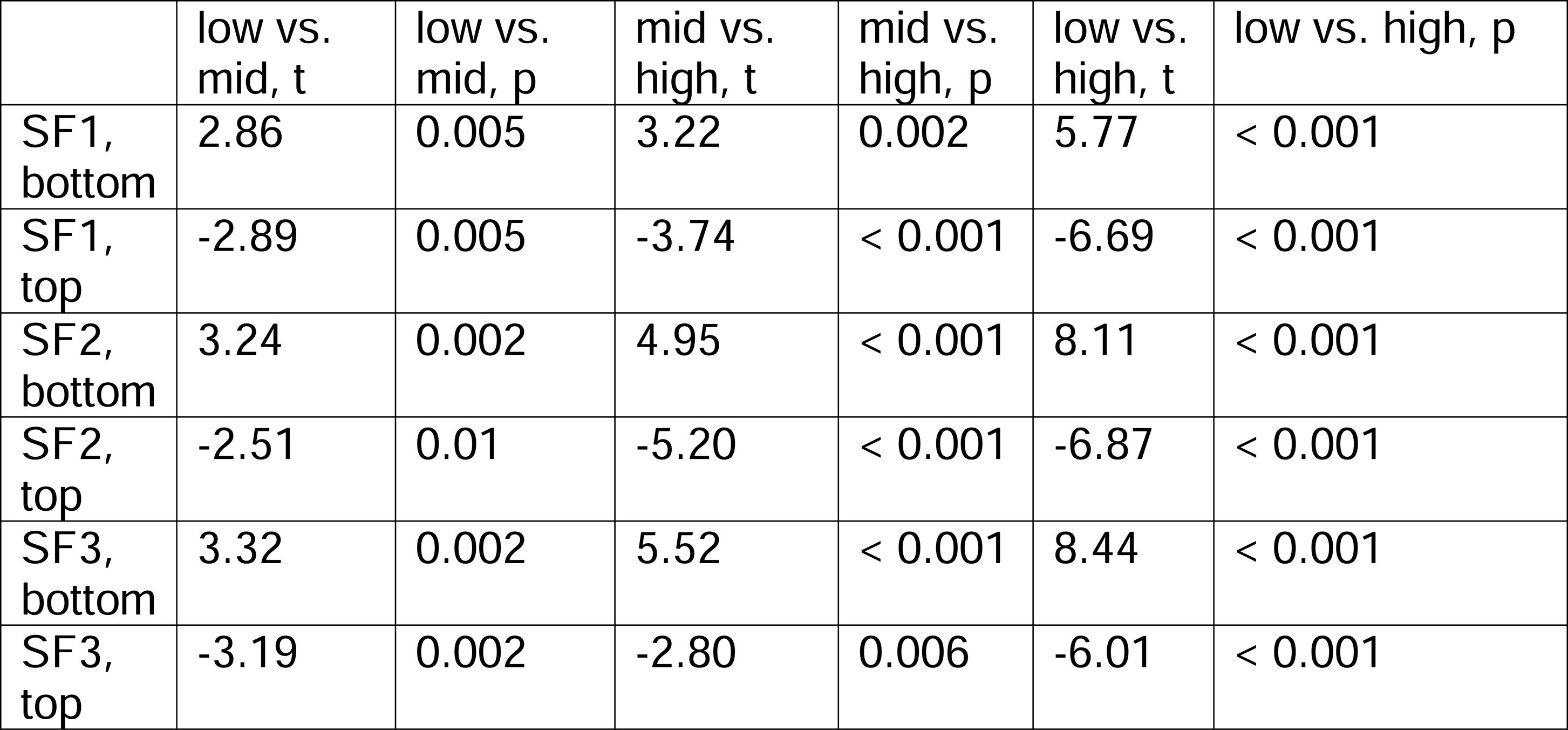
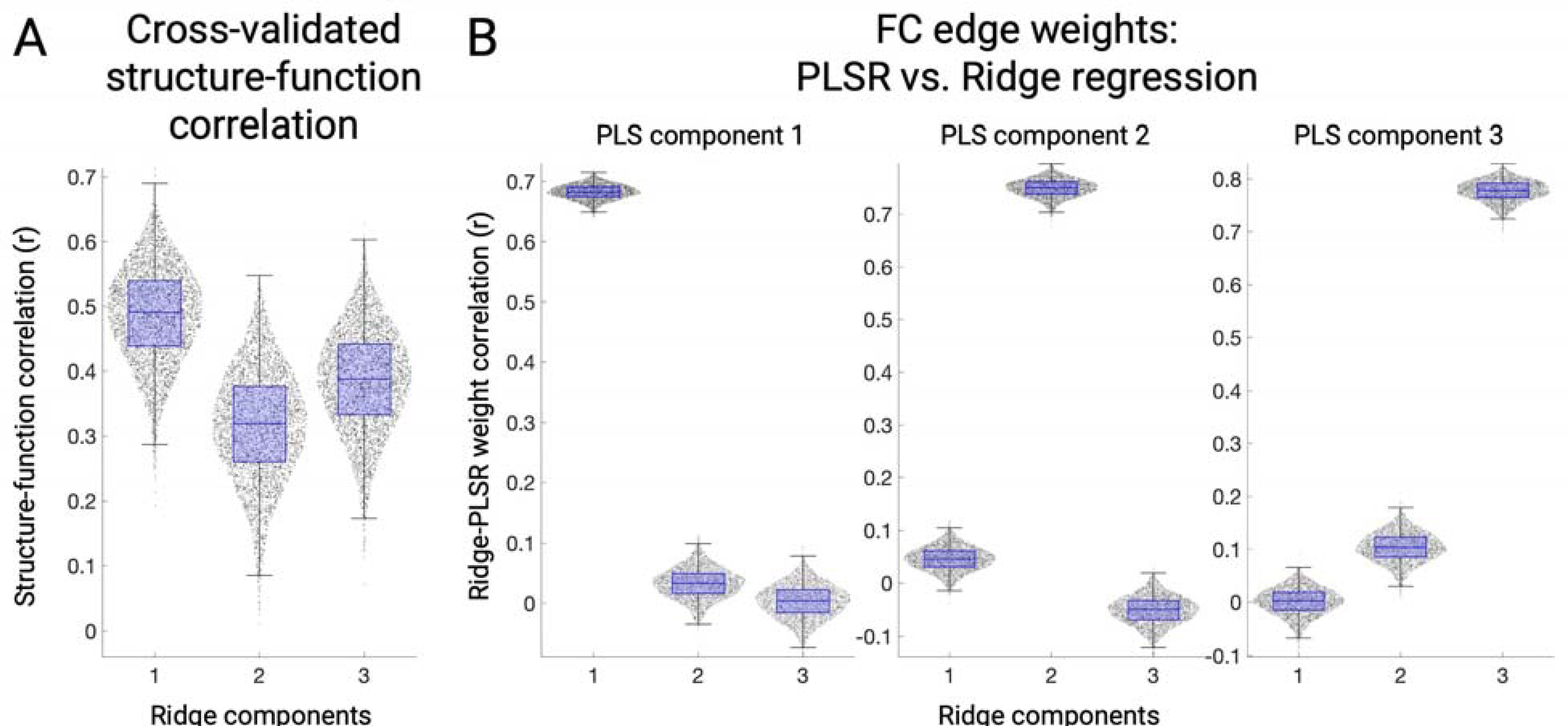
**A.** Out-of-sample correlation coefficients between atrophy component scores and ridge regression-derived functional connectivity scores. Each dot represents a single fold out of four folds per 1000 trials. **B.** Correlation coefficients between ridge regression functional connectivity edge weights (a [30135 x 1] vector) derived separately for each cross-validation fold versus partial least square regression-derived weights. The median correlations between corresponding ridge regression and PLSR components were 1-1: r=0.68, 2-2: r=0.75, 3-3: r=0.78.

**Supplementary Figure 4.**
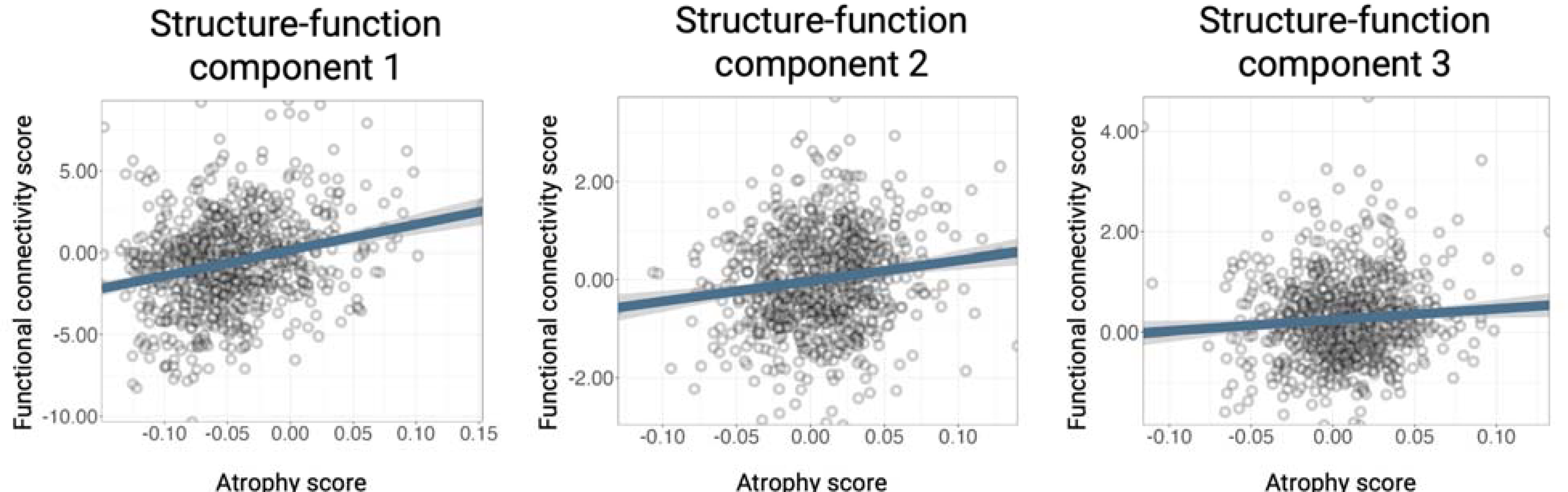
Structure-function component score correlations for the ADNI replication dataset. The partial correlation coefficients were SF1: r=0.25, p < 0.001, SF2: r=0.15, p < 0.001, SF3: r=0.08, p=0.015.

**Supplementary Figure 5.**
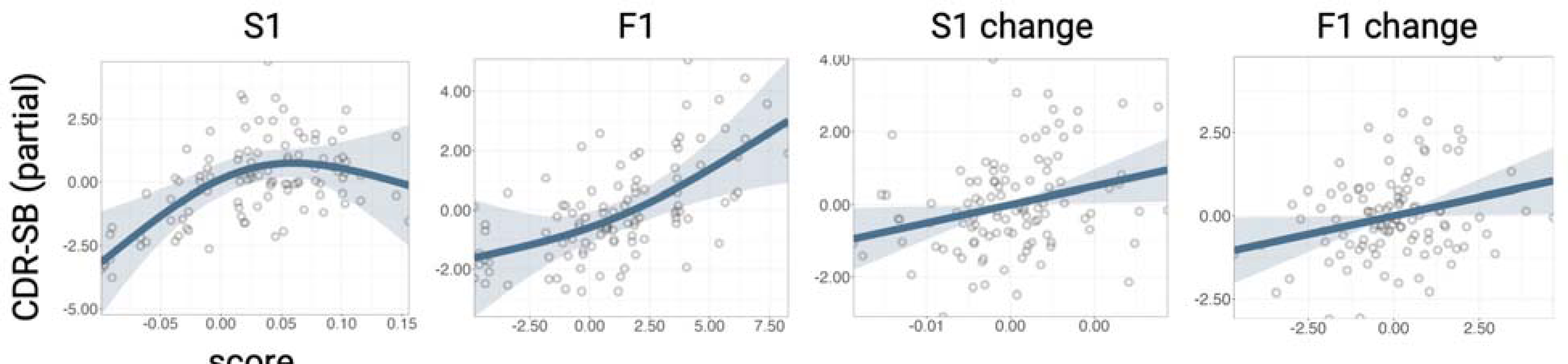
Longitudinal relationship effect plots showing the partial relationship of CDR-SB with structure component 1 (S1) mean (between-subject), function component 1 (F1) mean (between-subject), S1 change (within-subject), and F1 change (within-subject).

**Supplementary Figure 6.**
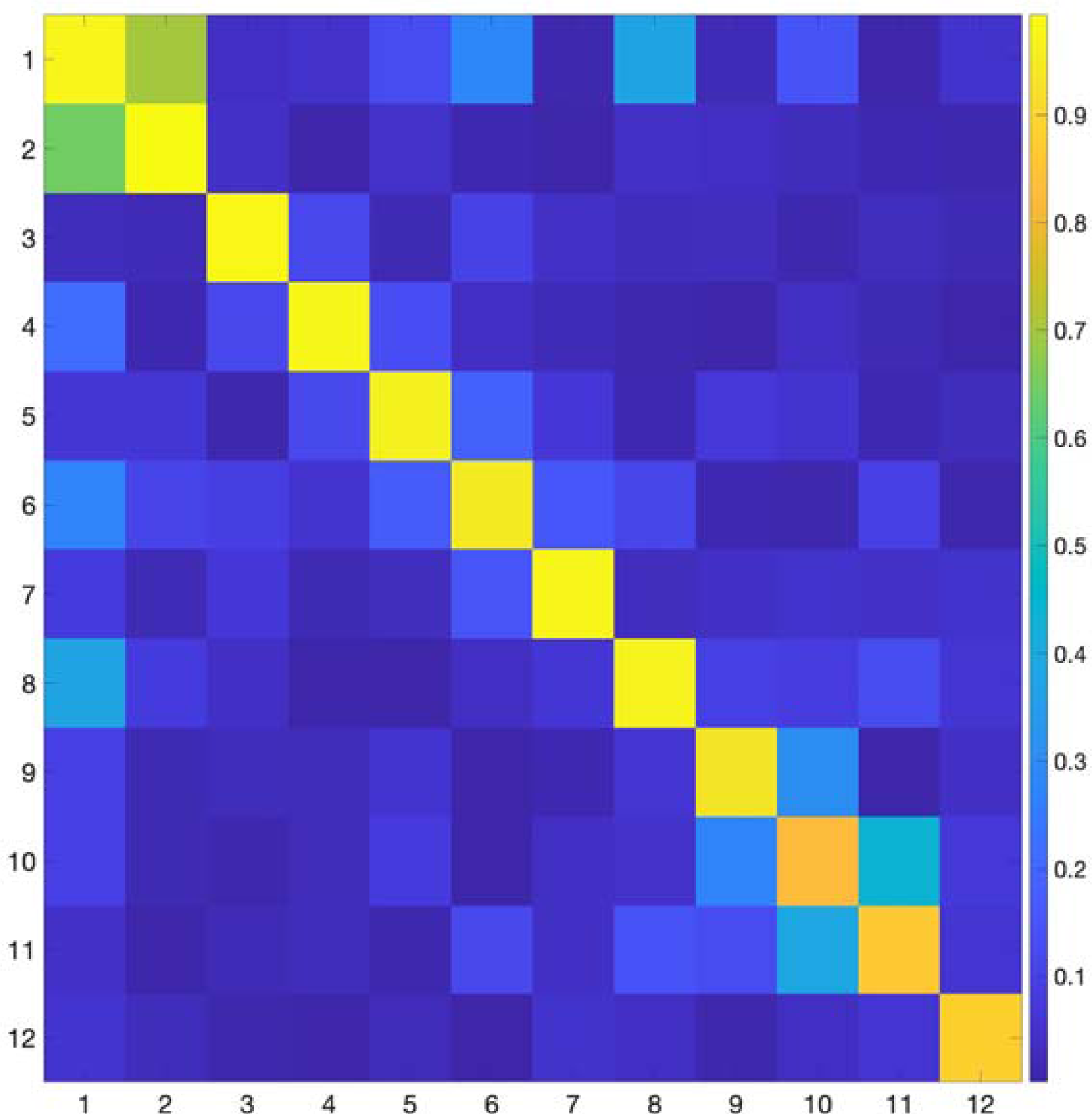
Spatial correlations between the fMRI PCA spatial components (gradients; n=246 regions per component) derived from the independent cognitively normal cohort (n=321, rows) and the main combined patient and control cohort (n=321, columns). **Supplementary Results 6.** The spatial gradient patterns were highly consistent in the independent cognitively normal cohort and the primary cohort. The median spatial correlation was r=0.98 for gradients 1-6 (max=0.99, min=0.95) and r=0.97 for gradients 1-12 (max=0.99, min=0.83).

**Supplementary Figure 7.**
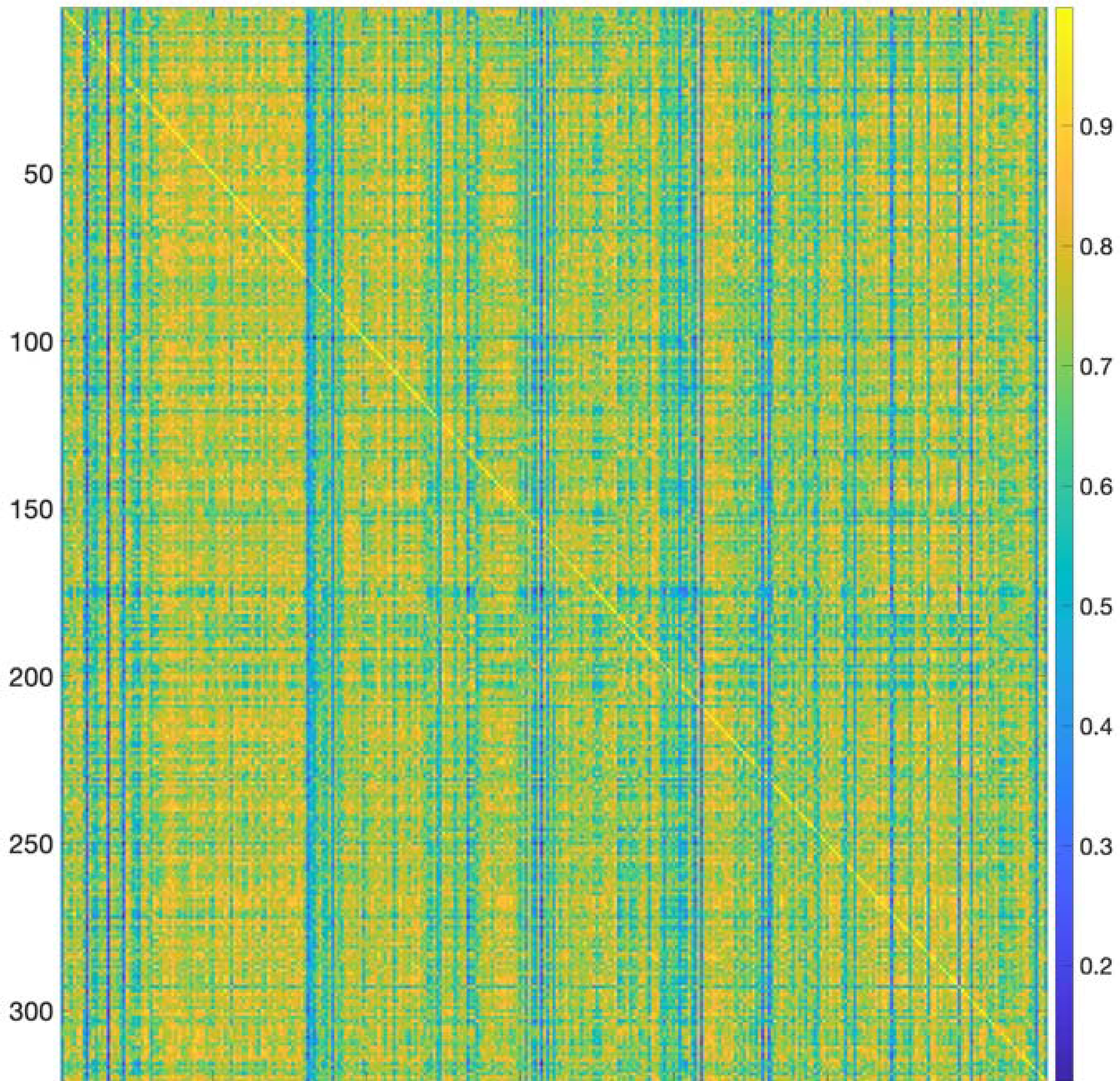
Correlations between each subject’s actual [246 x 246] FC matrix (rows) and simulated FC matrix (columns).

**Supplementary Figure 8.**
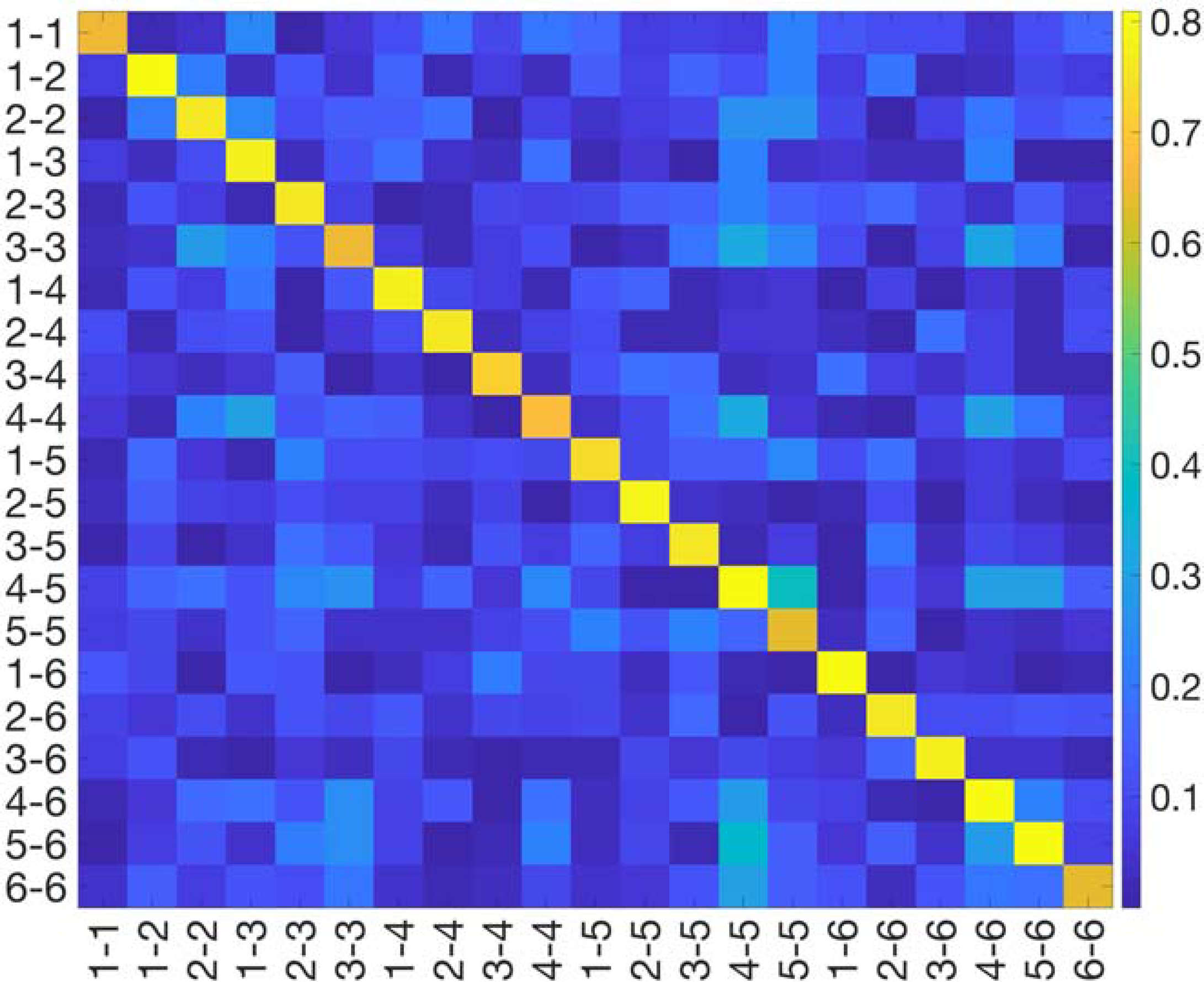
Correlations between observed gradient variance/covariance and coupling parameter-derived gradient amplitude/angles.

## Notes

### Competing Interest Statement

J.A.B. is a founder of Radiata Inc. J.A.B. has a financial interest in Radiata Inc. and may benefit financially if the company is successful in marketing biomarker software products.

### Summary of Updates

Modified the title and text in the abstract and introduction.

