## Supplementary Info for "Functional network collapse in neurodegenerative disease"

**Supplementary Information**

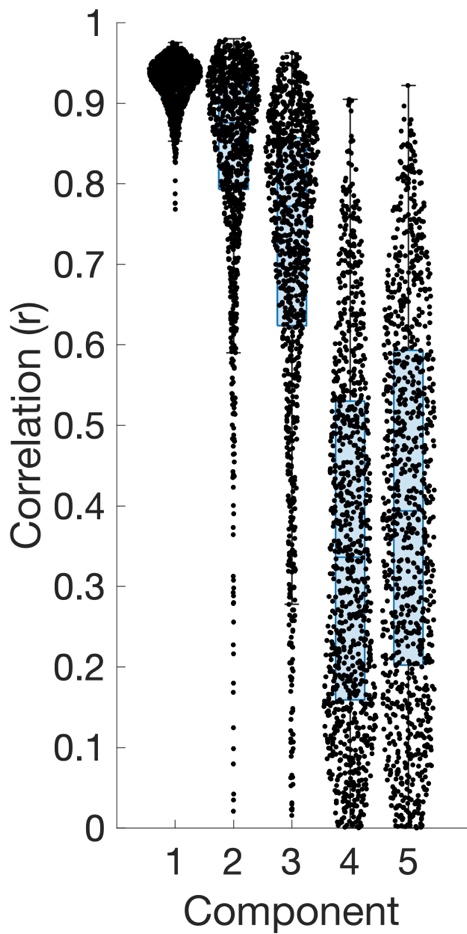

**Supplementary Figure 1.** Partial least squares regression structure component reliability using split-half analysis.

**Supplementary Results 1.** 1000 trials of independent PLSR were run on the first and second halves of the subjects, randomly split with balanced syndrome classes for each trial. The structure component loading vectors were correlated between split halves for the 1000 trials for the first five PLSR components. The median correlations were S1: r=0.93±0.03, S2: r=0.88±0.14, S3: r=0.77±0.20, S4: r=0.34±0.23, S5: r=0.39±0.24. The most substantial drop in reliability was between components 3 and 4.

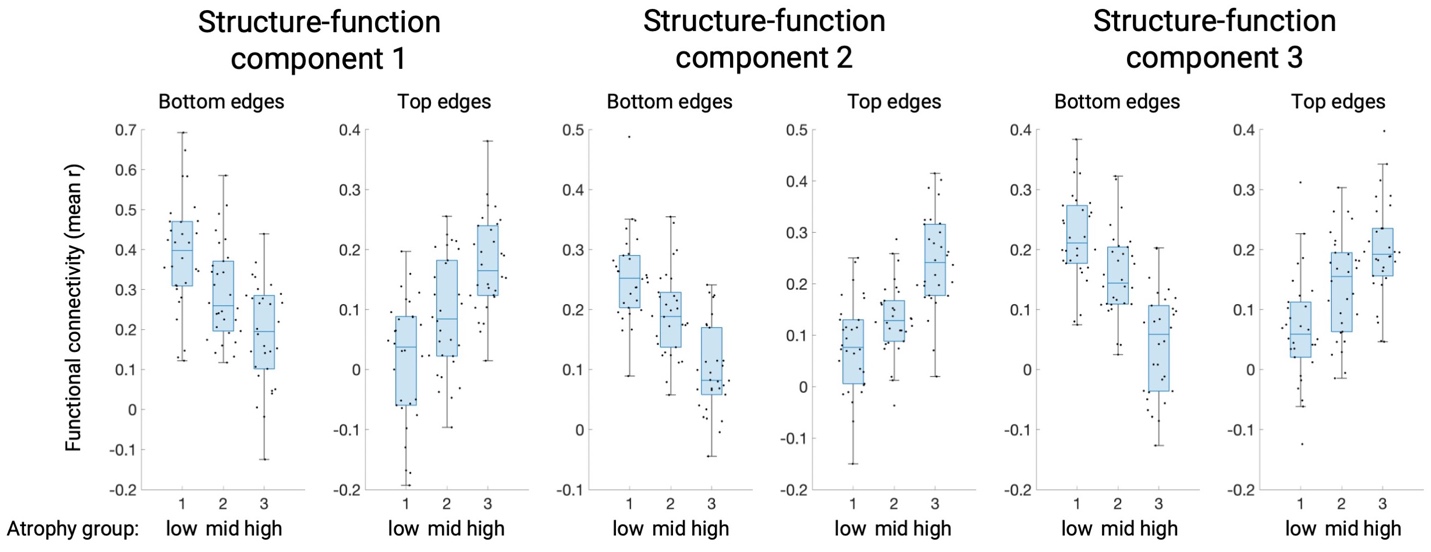

**Supplementary Figure 2.** FC edge weights for the bottom/top 1% of edges on each function component. Mean FC edge weights are shown for each component for groups of 30 subjects with the lowest/intermediate/highest atrophy scores. All boxplots show the median, lower and upper quartile range, and the non-outlier minimum/maximum.

**Supplementary Results 2.** For the first three structure-function components, subjects were sorted based on their structural score for that component and binned into groups of 30. For a given component, FC edges were sorted based on their PLSR weight. The bottom/top 1% of edges (300 edges) were kept and the mean FC weight for these two sets were computed for each subject. These mean edge weights were statistically compared for the groups of 30 subjects with low/middle/high atrophy on that component, both for the bottom and top FC edges. The edge weights were always statistically significant between low and high subjects (all p < 0.001, see table) and were significant at a less stringent threshold (p < 0.01) in all other tests. This indicated that the partial FC variance captured by the FC scores was substantial enough to capture significant differences in overall FC edge weights between groups of subjects.

|  | low vs. mid, t | low vs. mid, p | mid vs. high, t | mid vs. high, p | low vs. high, t | low vs. high, p |
| --- | --- | --- | --- | --- | --- | --- |
| SF1, bottom | 2.86 | 0.005 | 3.22 | 0.002 | 5.77 | < 0.001 |
| SF1, top | -2.89 | 0.005 | -3.74 | < 0.001 | -6.69 | < 0.001 |
| SF2, bottom | 3.24 | 0.002 | 4.95 | < 0.001 | 8.11 | < 0.001 |
| SF2, top | -2.51 | 0.01 | -5.20 | < 0.001 | -6.87 | < 0.001 |
| SF3, bottom | 3.32 | 0.002 | 5.52 | < 0.001 | 8.44 | < 0.001 |
| SF3, top | -3.19 | 0.002 | -2.80 | 0.006 | -6.01 | < 0.001 |

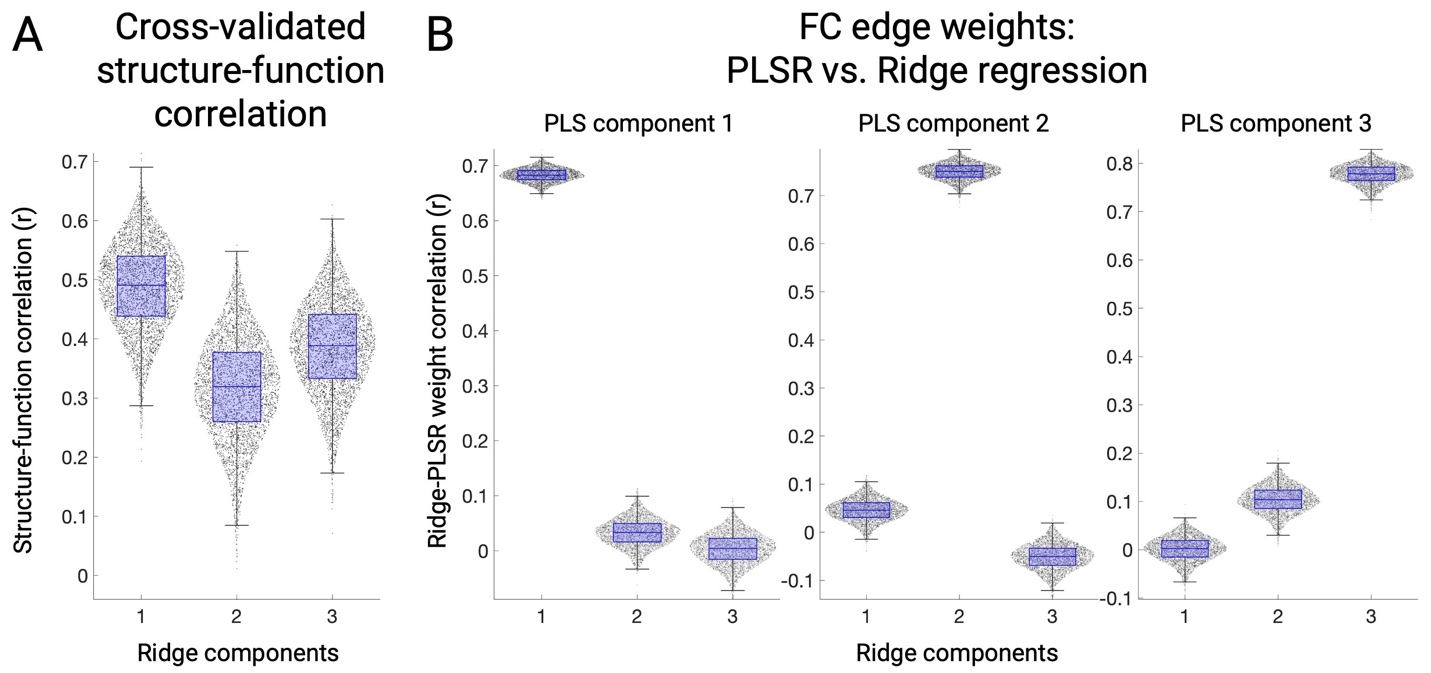

**Supplementary Figure 3. A.** Out-of-sample correlation coefficients between atrophy component scores and ridge regression-derived functional connectivity scores. Each dot represents a single fold out of four folds per 1000 trials. **B.** Correlation coefficients between ridge regression functional connectivity edge weights (a [30135 x 1] vector) derived separately for each cross-validation fold versus partial least square regression-derived weights. The median correlations between corresponding ridge regression and PLSR components were 1-1: r=0.68, 2-2: r=0.75, 3-3: r=0.78.

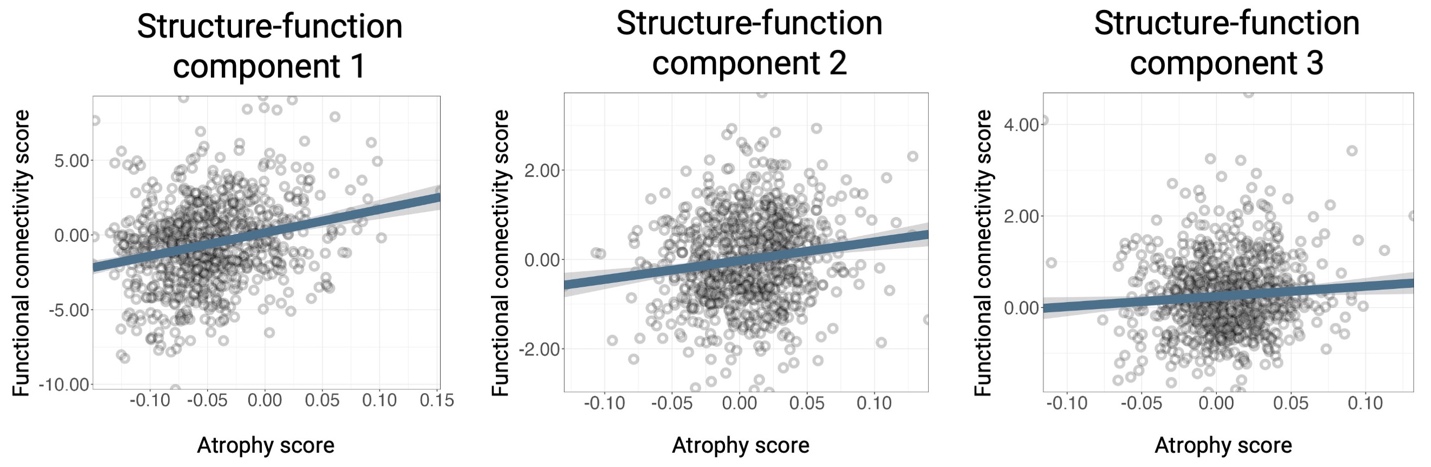

**Supplementary Figure 4.**

Structure-function component score correlations for the ADNI replication dataset. The partial correlation coefficients were SF1: r=0.25, p < 0.001, SF2: r=0.15, p < 0.001, SF3: r=0.08, p=0.015.

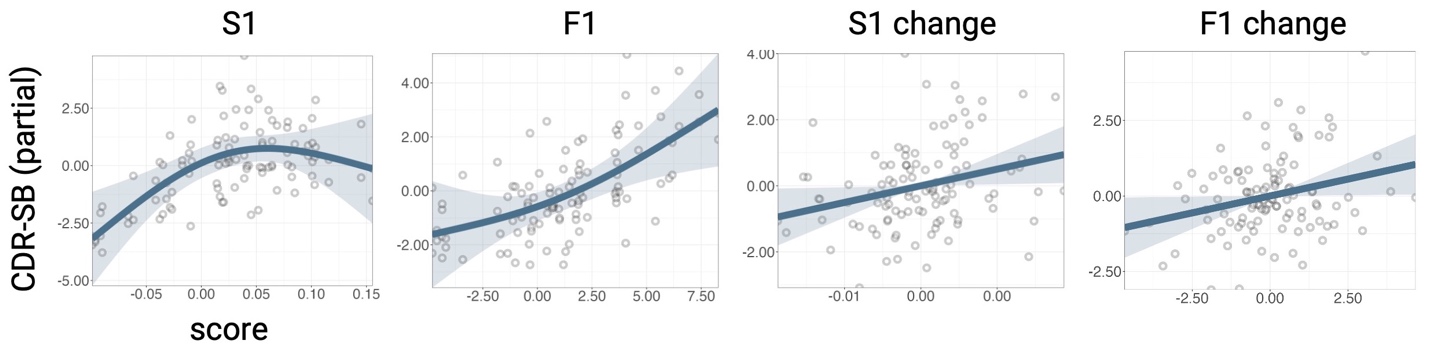

**Supplementary Figure 5.** Longitudinal relationship effect plots showing the partial relationship of CDR-SB with structure component 1 (S1) mean (between-subject), function component 1 (F1) mean (between-subject), S1 change (within-subject), and F1 change (within-subject).

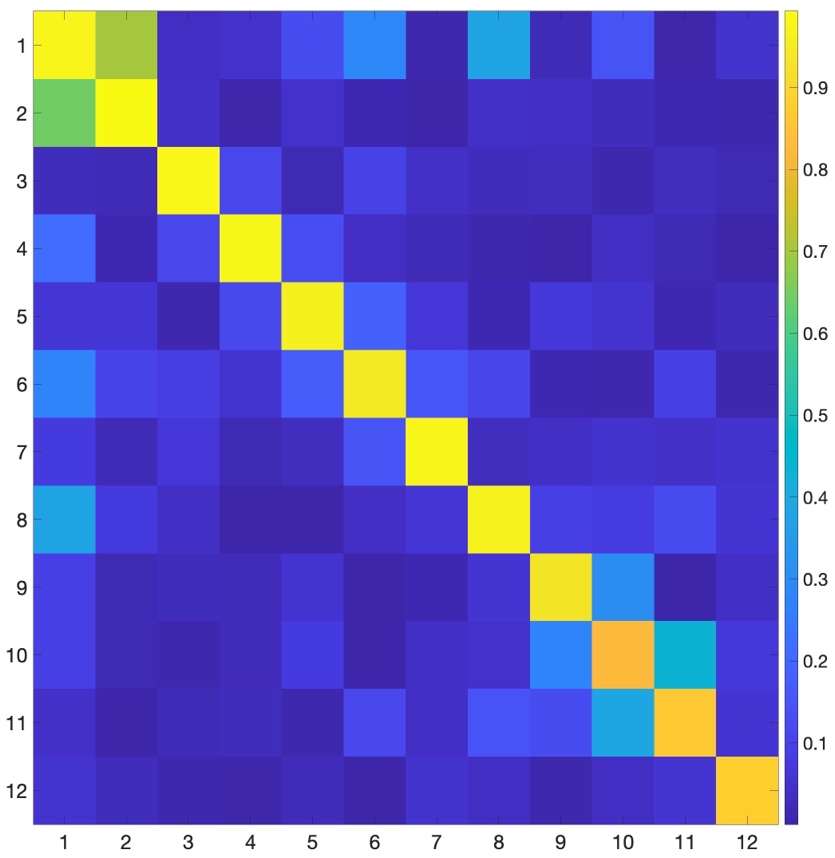

**Supplementary Figure 6.** Spatial correlations between the fMRI PCA spatial components (gradients; n=246 regions per component) derived from the independent cognitively normal cohort (n=321, rows) and the main combined patient and control cohort (n=321, columns).

**Supplementary Results 6.**

The spatial gradient patterns were highly consistent in the independent cognitively normal cohort and the primary cohort. The median spatial correlation was r=0.98 for gradients 1-6 (max=0.99, min=0.95) and r=0.97 for gradients 1-12 (max=0.99, min=0.83).

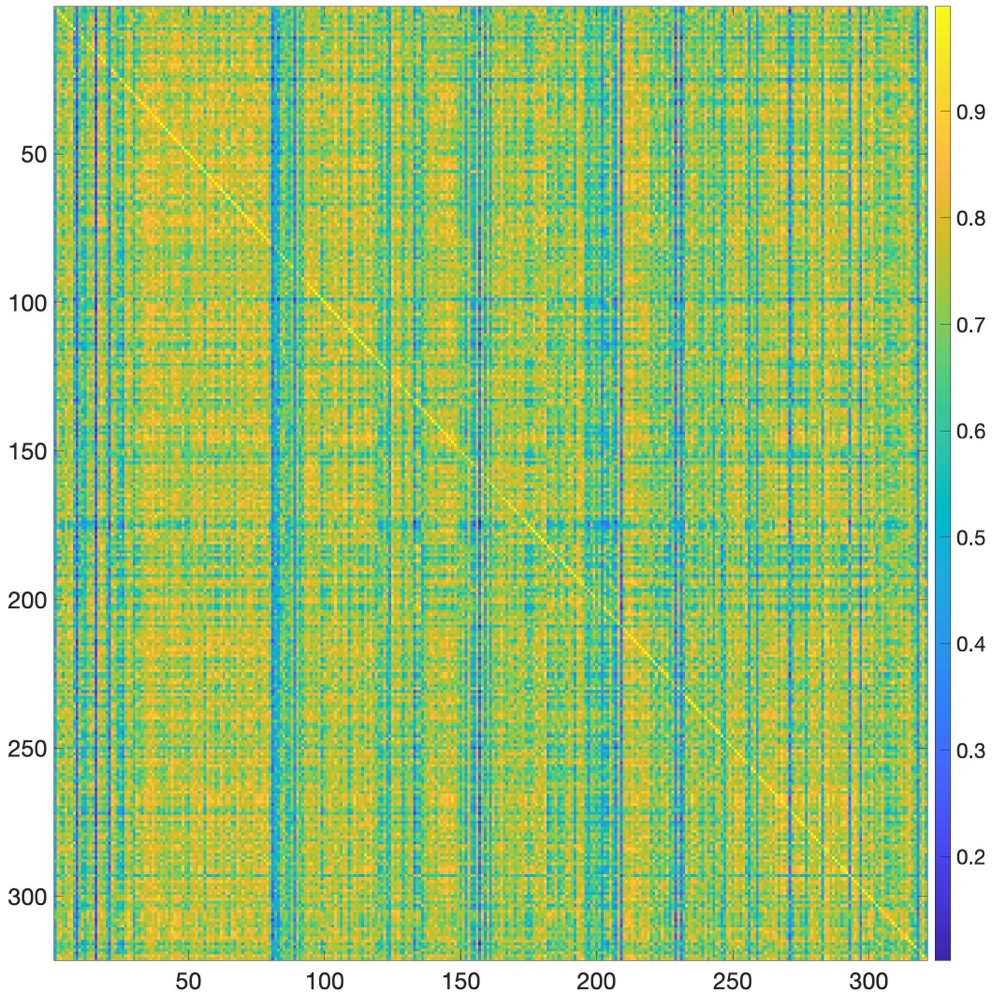

**Supplementary Figure 7.** Correlations between each subject's actual [246 x 246] FC matrix (rows) and simulated FC matrix (columns).

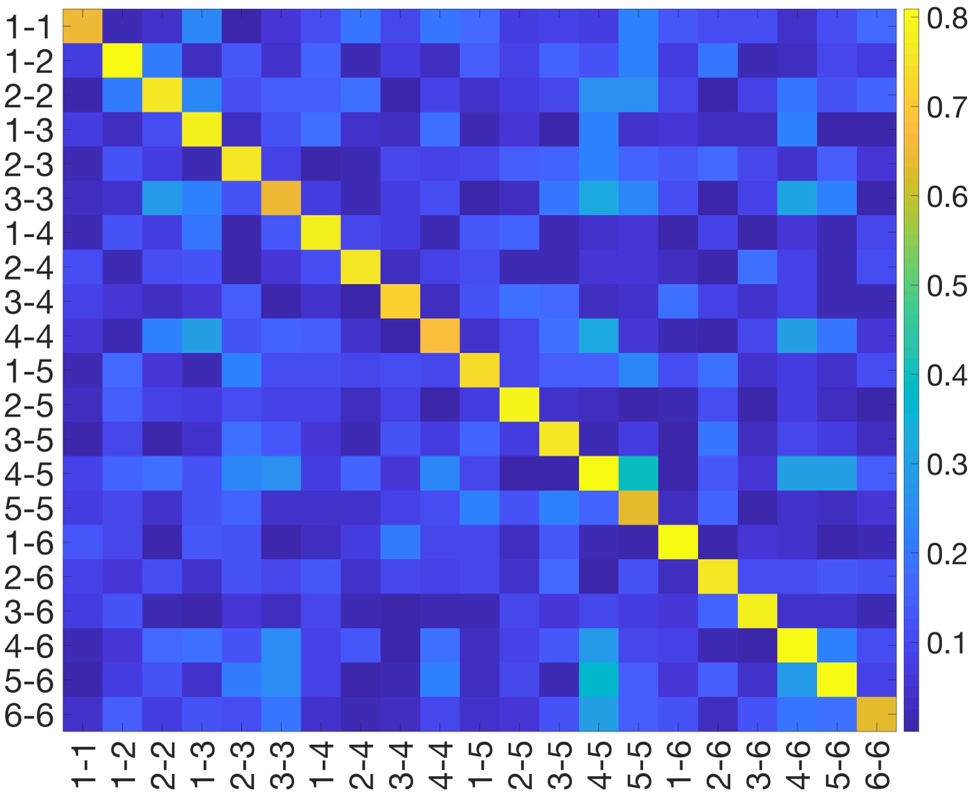

**Supplementary Figure 8.** Correlations between observed gradient variance/covariance and coupling parameter-derived gradient amplitude/angles.
